# Detection of H3N8 influenza A virus with multiple mammalian-adaptive mutations in a rescued Grey seal (*Halichoerus grypus*) pup

**DOI:** 10.1101/741173

**Authors:** Divya Venkatesh, Carlo Bianco, Alejandro Núñez, Rachael Collins, Darryl Thorpe, Scott M. Reid, Sharon M. Brookes, Steve Essen, Natalie McGinn, James Seekings, Jayne Cooper, Ian H. Brown, Nicola S. Lewis

## Abstract

Avian Influenza A Viruses (IAV) in different species of seals display a spectrum of pathogenicity, from subclinical infection to mass mortality events. Here we present an investigation of avian IAV infection in a 3-4 month old Grey seal (*Halichoerus grypus*) pup, rescued from St Michael’s Mount, Cornwall in 2017. The pup underwent medical treatment but died after two weeks; post-mortem examination and histology indicated sepsis as the cause of death. IAV NP antigen was detected by immunohistochemistry in the nasal mucosa, and sensitive real-time reverse transcription polymerase chain reaction assays detected trace amounts of viral RNA within the lower respiratory tract, suggesting that the infection may have been cleared naturally. IAV prevalence among Grey seals may therefore be underestimated. Moreover, contact with humans during the rescue raised concerns about potential zoonotic risk. Nucleotide sequencing revealed the virus to be of subtype H3N8. Combining a GISAID database BLAST search and time-scaled phylogenetic analyses, we inferred that the seal virus originated from an unsampled, locally circulating (in Northern Europe) viruses, likely from wild Anseriformes. From examining the protein alignments, we found several residue changes in the seal virus that did not occur in the bird viruses, including D701N in the PB2 segment, a rare mutation, and a hallmark of mammalian adaptation of bird viruses. IAVs of H3N8 subtype have been noted for their particular ability to cross the species barrier and cause productive infections, including historical records suggesting that they may have caused the 1889 pandemic. Therefore, infections such as the one we report here may be of interest to pandemic surveillance and risk and may help us better understand the determinants and drivers of mammalian adaptation in influenza.

## INTRODUCTION

Influenza A viruses (IAVs) are important pathogens for humans and livestock including pigs and poultry. They are segmented RNA viruses, whose genomes consist of eight segments of RNA, which code for ~11 proteins/polypeptides. IAVs are classified into several subtypes based on the antigenic properties of two surface glycoproteins haemagglutinin (HA, avian subtypes H1-H16) and neuraminidase (NA, avian subtypes N1-N9). Viruses of most subtypes can be found in wild waterfowl and shorebirds which are their natural reservoir (Alexander, 2007; Easterday et al., 1968) and can infect both domestic birds and mammalian species in spill-over infections. A few IAV lineages have become established in mammals such as humans, pigs, horses, and dogs though only of subtypes H1, H2 and H3 in combination with N1, N2 or N8 (Reperant et al., 2009; Webster et al., 1992).

IAV in birds replicates mainly in the intestine and is transmitted through the faecal-oral route, although respiratory tropism and oropharyngeal shedding has been noted (Webster et al., 1978; Daoust et al., 2011; Höfle et al., 2012; França et al., 2012; Daoust et al., 2013), and the virus can survive in the environment for fairly long periods (Brown et al., 2009; Stallknecht et al., 1990). This creates conditions conducive to viral exchange with marine mammals such as pinnipeds (seals), whose habitats and prey intersect with those of waterfowl and shorebirds. Several cases of marine mammal infection with IAV of many subtypes including H1N1, H3N3, H3N8, H4N5, H4N6, H7N7 and H10N7 have been documented with a spectrum of effects ranging from mass die-offs to asymptomatic; in a majority of these cases, the source is implicated to be avian (Fereidouni et al., 2016; White, 2013). A few studies have also suggested that Grey seals may act as an endemically infected reservoir which may disseminate viruses in coastal ecosystems to other mammals, coastal birds, and potentially humans (Duignan et al., 1995, 1997; Puryear et al., 2016). Indeed, seroprevalence levels in live-captured healthy Grey seal populations (20 – 26%) (Bodewes et al., 2015; Puryear et al., 2016) is comparable with levels found in wild birds (31% - 60%) depending on species, geography, seasonality and other factors (Curran et al., 2015; Fereidouni et al., 2010; Wilson et al., 2013).

In seals, IAV binds the same type of sialyloligosaccharide receptors as birds (SAα2,3Gal), but the receptors are located in the respiratory tract and lungs instead of the intestinal tract, suggesting that inhalation is the most likely route of transmission (Ito et al., 1999; Ramis et al., 2012; White, 2013). Seals may also be infected by IAV from non-avian sources; there is serological evidence for infection of Baikal and ringed seals in Russia with human H3N2 strains A/Aichi/2/68 and A/Bangkok/1/79 (Ohishi et al., 2004) and a human pandemic-2009 H1N1 virus was isolated from Elephant seals in 2010 on the coast of California, USA. Furthermore, it has been shown that IAV from seals can replicate in human tissue and that seal IAV can be systemically virulent in primates (Murphy et al., 1983; Webster et al., 1981; White, 2013). Seals may therefore be potential sources of pandemic influenza.

In this paper, we report an H3N8 IAV infection of a rescued Grey seal pup in coastal England. We provide molecular and histological evidence for presence of IAV in the seal’s respiratory tissues, and conclude from the post-mortem that the clinical presentation was not caused by IAV. To our knowledge, we provide the first whole genome sequence for an IAV isolated from a Grey seal, as attempts so far have been unsuccessful. We use phylogenetic analyses to find the putative sources of this virus and look for adaptive changes in its sequence. We compare this case with previously described seal infections to identify unique or parallel elements which may have implications for animal and human health.

## METHODS

### Clinical history and medical interventions

In February 2017, a female Grey seal (*Halichoerus grypus*) pup was rescued from St Michael’s Mount, Cornwall. Physical examination of the animal on admittance to the rehabilitation centre revealed an emaciated and dehydrated subject (body weight = 20 kg). The estimated age was 3 to 4 months. The animal exhibited a mucoid nasal discharge and a 1 cm wound on the ventral thorax. Its temperature was 39.9 ^o^C and breathing rate was abnormal (continuous breathing pattern, 12 breaths per minute). The pup was monitored and provided nutritional support, fluid therapy and antibiotic treatment, but died suddenly 14 days after admittance. The carcass and a nasal swab were submitted to the Animal and Plant Health Agency (APHA) for virological investigation.

### Pathology, histopathology and immunohistochemistry

The carcass underwent a full post-mortem examination. A set of tissues was sampled and fixed in buffered formalin (nasal turbinates, trachea, lung, kidney, soft tissues adjacent to the cutaneous/subcutaneous lesion, and lymph node). A standard histopathological examination was carried out on the tissues (Hematoxylin & eosin), followed by an immunohistochemical investigation targeting nucleoprotein to detect intra-lesional IAV along the respiratory tract (Brookes et al., 2010).

### Real-time reverse transcription polymerase chain reaction (RRT-PCR)

RNA was extracted from the nasal swab and tissue suspensions using the QIAmp viral RNA BioRobot kit customised for APHA in conjunction with a Universal BioRobot (Qiagen, Manchester, UK) (Slomka et al., 2009). RRT-PCR testing of the RNA extracts comprised (i) the Matrix (M)-gene assay for generic IAV detection using the primers and probes of (Nagy et al., 2010) and (ii) H5 and H7 IAV RRT-PCR assays to test for notifiable avian influenza (Slomka et al., 2007, 2009). For each RRT-PCR assay, samples producing a threshold cycle (CT) value <36.0 were considered positive (Slomka et al., 2010). The RNA was also tested by an IAV N1-specific RRT-PCR according to the procedure described by (Payungporn et al., 2006), an IAV N5-specific RRT-PCR (James et al. 2018) and two IAV subtype N8-specific RRT-PCRs (James et al., 2018) with the same positive/negative acceptance criteria. All amplifications were carried out in an MX3000P qPCR System (Agilent).

### Virus isolation and whole genome sequencing

Attempted virus isolation in 9-to 11-day old SPF embryonated fowls’ eggs was performed on the nasal swab sample according to the internationally-recognised European Union (EU) and OIE methods (EU, 2006; OIE, 2015), but was unsuccessful. RNA was sequenced using the MiSeq platform. Briefly, viral RNA was extracted from the egg-isolate of the virus from the nasal swab using the QIAmp viral RNA mini-kit without the addition of carrier RNA (Qiagen, Manchester, UK). cDNA was synthesized from RNA using a random hexamer primer mix and cDNA Synthesis System (Roche, UK). The Sequence library was prepared using a NexteraXT kit (Illumina, Cambridge, UK). Quality control and quantification of the cDNA and Sequence Library was performed using Quantifluor dsDNA System (Promega, UK). Sequence libraries were run on a Miseq using MiSeq V2 300 cycle kit (Illumina, Cambridge, UK) with 2×150 base paired-end reads. The raw sequence reads were analysed using publicly available bioinformatics software, following an in house pipeline, available on github (https://github.com/ellisrichardj/FluSeqID/blob/master/FluSeqID.sh). This pipeline de-novo assembles the raw data using the Velvet assembler (Zerbino and Birney, 2008), blasts the resulting contigs against a local database of influenza genes using Blast+ (Camacho et al., 2009), then maps the raw data against the highest scoring blast hit using the Burrows-Wheeler Aligner (Li, 2013). The consensus sequence was extracted from the resultant bam file using a modified SAMtools software package (Li et al., 2009), script (vcf2consensus.pl) available at: https://github.com/ellisrichardj/csu_scripts/blob/master/vcf2consensus.pl).

### Phylogenetic analysis of the seal virus

BLAST (Basic Local Alignment Search Tool) was used on GISAID (Elbe and Buckland-Merrett, 2017; Shu and McCauley, 2017) to find the top 50 closest related viral segments for each segment of the seal virus. We combined the seal virus sequence (query) along with the BLAST-hits sequences (blasthit) for each segment for phylogenetic analysis. We removed sequences containing duplicate strain names and aligned with MAFFT (Katoh and Standley, 2013) using automatic settings. Alignments for each segment were inspected manually on AliView (Larsson, 2014) and the ends trimmed to the starting ATG and end STOP codon. Exploratory trees were run using FastTree (Price et al., 2009), after which we used IQ-TREE (Nguyen et al., 2015) to make the final maximum-likelihood tree with 1000 iterations of alrt (approximate likelihood ratio test) for branch support. Tempest v1.5 (Rambaut et al., 2016) was used to test ML trees for clock-like behaviour. Trees for all segments except MP showed clock-like behaviour (**Figure S1**), so results from the MP dataset were excluded. BEAST v1.10.1 (Bayesian Evolutionary Analysis Sampling Trees) (Suchard et al., 2018) was used to determine the putative time and source of emergence of the different segments of the seal virus. BEAST performs Bayesian analysis of molecular sequences using MCMC (Markov Chain Monte Carlo) methods. For all segments other than MP and NS which have multiple reading frames, we used the SRD06 site model which partitions the codons into 1+2 and 3. For NS we used a GTR model with no codon partitioning. The rest of the priors were kept identical for all segments: an uncorrelated relaxed lognormal clock, GMRF (Gaussian Markov random field) Bayesian skyride tree (Minin et al., 2008), 7 million MCMC generations with sampling every 7000. Two separate runs were performed for each segment, which were combined after logs were inspected in Tracer v1.7.1(Rambaut et al., 2018) for appropriate mixing and ESS (effective sample size) values > 200. Trees were summarised into median clade credibility trees (MCC) and plotted in R v3.5 (using the ggtree package (Yu et al., 2017).

### Amino acid substitutions

Trimmed alignments of each segment were manually inspected in AliView software (Larsson, 2014), translated into amino-acids, and checked for amino acid changes across each dataset. We identified several substitutions in the seal virus that did not occur in any of the bird virus sequences. We recorded these substitutions, and used the H3-numbering of the sequence using the HA subtype numbering conversion tool available from FluDB ((Burke and Smith, 2014), https://tinyurl.com/HAnumbering) for the HA protein. We also looked for differences in glycosylation patterns between the seal and the related wild bird HA and NA glycoproteins using a program to detect Asn-X-Ser or Asn-X-Thr (where X is any amino acid other than Proline) patterns in the amino acid sequences (personal communication from Todd Davis, CDC, USA)

## RESULTS

### Pathology and immunohistochemistry

The post-mortem revealed abundant purulent fluid (approximately 1 litre) in the subcutaneous tissues of the right side of the body extending from the neck to the thoracic region (Figure 1A). All other organs were macroscopically unremarkable. The blubber sternal thickness was 0.4 cm. Histology identified a severe, chronic-active, fibrino-purulent cellulitis and fasciitis and a severe, acute renal infarct with thrombosis. In the respiratory system a diffuse mild to moderate rhinitis with epithelial hyperplasia and presence of mites was observed in the nasal cavity and a focal broncho-interstitial pneumonia with thrombosis and pulmonary nematodes were seen in the lungs, with no changes observed in trachea and bronchi. Influenza A virus antigen was detected by immunohistochemistry only in the nasal mucosa, in the nuclei of scattered isolated epithelial cells (Figure 1B). The pathological findings were suggestive of a thromboembolic event and sepsis caused by the cellulitis as the cause of the death of the animal.

**Figure 1 (A-B).**
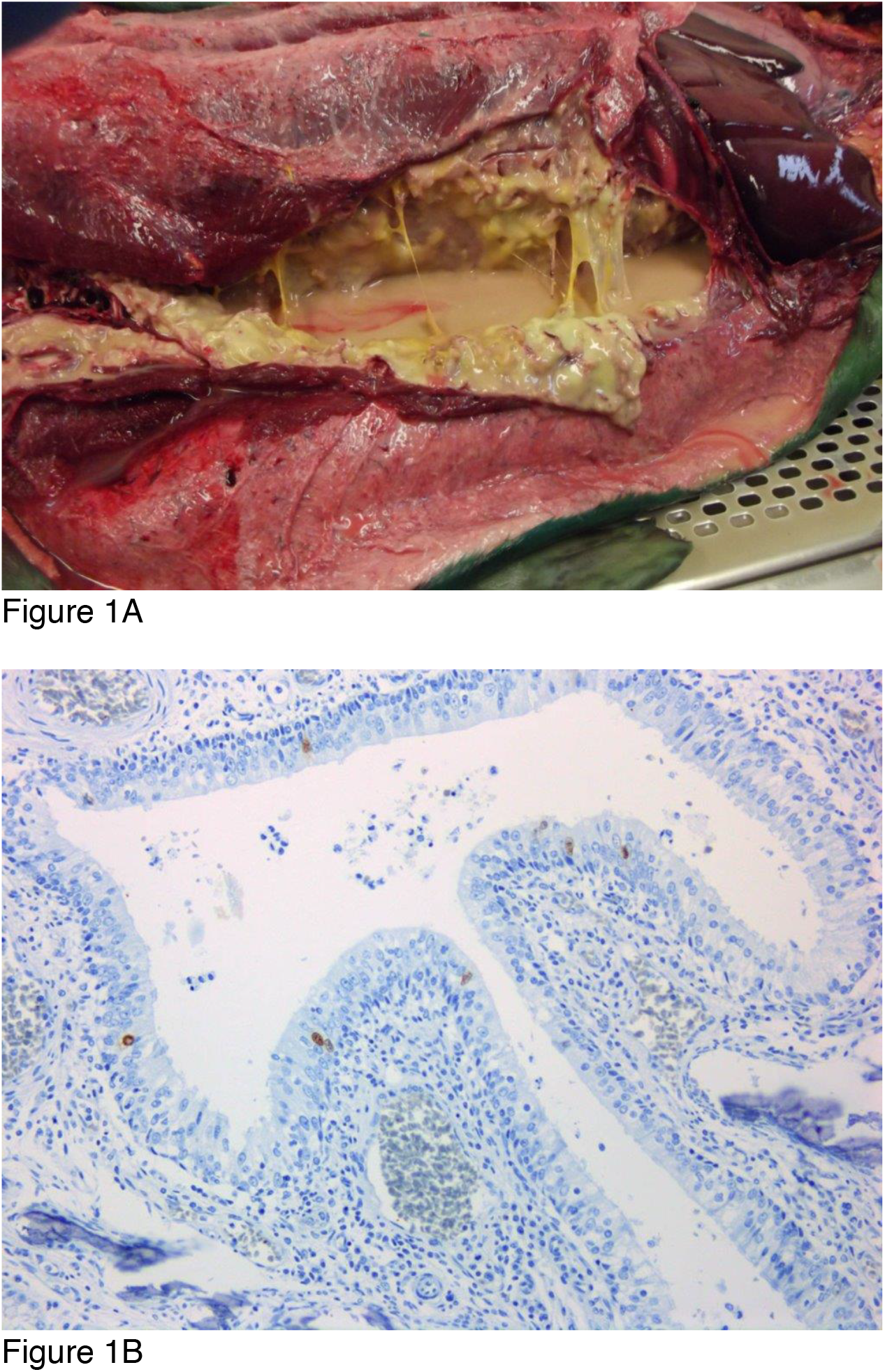
Grey seal pup. (A) Focally extensive, chronic-active, purulent and fibrino-necrotising cellulitis in subcutaneous tissues and skeletal muscle over the thorax. (B) Immunohistochemical demonstration of Influenza A nucleoprotein antigen in the nuclei of epithelial cells within the nasal mucosa of a grey seal. (DAB chromogen and haematoxylin counterstain) (Objective: 40x).

### Detection and subtyping of IAV by RRT-PCR

The nasal swab was positive for the M-gene (CT value of 22.36), signifying the presence of IAV RNA but was negative for NAI haemagglutinin (HA) subtypes H5 and H7 as well as the IAV subtypes N1 and N5. However, IAV subtype N8 was detected by one of the specific RRT-PCR assays employed (CT value of 24.38). In addition, trace amounts of IAV RNA were detected by the M-gene RRT-PCR assay in tracheal bronchi (CT value of 35.88), the left lung (CT value of 37.73) and the right cranial lung (CT value of 38.67). Due to low levels of virus detected in the lower respiratory tract, and the presence of clinical signs not consistent with influenza infection, we concluded that the virus was of no or low pathogenicity. It is possible that at the point the seal was rescued, it had already cleared the infection in the lungs naturally, which would explain trace amounts of detection in these tissues. This is consistent with previous studies which found that Grey seals were likely to remain asymptomatic with IAV infection while a proportion of the surveyed adults and juveniles were seropositive (Bodewes et al., 2015; Puryear et al., 2016).

Sequencing results from the nasal swab egg-isolate indicated no mixed infection, just a single virus, from which it was possible to sequence all eight gene segments. The virus-derived sequence was named A/Grey seal/England/027661/2017 and sequences were deposited in GISAID database (Global Initiative on Sharing All Influenza Data) with isolate ID: EPI_ISL_381748.

### Source of viral segments

Segment gene sequences which were found to be most closely related to the genes of the seal virus by BLAST were of avian origin. Overall, the strains in the set of BLAST-hits for each segment gene, all came from different avian viruses, isolated from different bird-types, and of different subtypes. They were from strains that were mostly isolated from wild birds such as mallard, Eurasian teal, White-fronted goose, Black-headed gull (*Anas platyrhynchos*, *Anas crecca*, *Anser albifrons*, *Chroicocephalus ridibundus*) and others, along with a few from domestic birds such as chickens (*Gallus gallus domesticus*). This indicates a wild bird origin for the seal virus, which is consistent with previously detected seal infections (Fereidouni et al., 2016; White, 2013), and the hypothesis that overlapping habitats for wild birds and seals make transmission from wild birds possible, but less likely from poultry, pigs or humans.

Also similar to previously reported infections (Anthony et al., 2012; Zohari et al., 2014), the set of closest related strains were isolated from within the local region, in this case, Northern Europe (The Netherlands, Germany, UK etc.) with the exception of three strains from China being among the close relatives of the PB1 sequence. The years of detection of the closest-related avian viruses for each segment ranged from the 2007 (NA) to 2017 (NS).

**Table S1** shows a summary for how often the same strain appears as a BLAST-hit for each segment of the seal virus. A large proportion of these occur singly (250 strains) or for a maximum of two segments (40 strains). However, some strains were found to map for 3, 4 or 5 segments (19, 2 and 1 strains(s) respectively). Of these, all the strains that mapped for at least 3 segments were isolated from birds in the Netherlands between 2011-15, except for one chicken virus from France in 2016. The subtypes varied greatly, but they were usually isolated from wild birds. In the next section, we further examine the closest available sequence(s) for each seal gene segment to try and understand the emergence and further propagation of this virus.

### Emergence of viral segments

There is no dedicated surveillance program for seals in England so infection status in seals as endemic or one-off spill over event is uncertain. Previous studies in the US have indicated that Grey seals are possibly a reservoir for IAV and other viruses (Duignan et al., 1995, 1997; Puryear et al., 2016), We performed a time-scaled analysis with the set of closely related avian virus segment sequences, to test if all segments from seal virus had a similar point of introduction, i.e., similar time to most recent common ancestor (TMRCA) of the seal and wild bird viruses for all segments, or the extent to which this varied segment to segment. We used BEAST to reconstruct time-scaled phylogenies for each of our segment datasets.

Maximum clade credibility trees for each segment are presented in Figure 2 (A-G); a maximum-likelihood tree is presented for the MP gene segment in Figure 2H as it was excluded from BEAST analysis due to lack of clock-like behaviour in the dataset. Time to the putative ancestor strain of the segments from the seal virus and its closest related segment is inferred as ranging from 1999 (NA) to between 2011 and 2015 (all other gene segments), summarised in Figure 3 and Table 1. Gaps in surveillance, and the availability of just one seal strain will likely affect the inference of TMRCAs, but the variation between segments is also likely a testament to the high levels of reassortment seen in wild bird IAVs (Lu et al., 2014). Indeed, the closest associated virus varies in host, subtype and geography of isolation for each segment, as can be seen in the highlighted clades in Figure 2 (A-H), and summarised in Table 1. Consistent with data of BLAST hits (Table S1), many of the closest related segments are from strains isolated in the Netherlands (HA, MP, NP, NS, PA, PB1, PB2). Where multiple strains are equally closely related to the seal virus, e.g. for PB1 and PB2, the avian influenza strains come from the Netherlands and France. The closest strain to the seal NA gene that has been sampled is from 2009 at the latest, from Norway. For the NS gene, the closest segments appear to be from Russian strains in chickens as recent as December 2017, which may indicate onward transmission via unsampled intermediaries; however the posterior probability support value for the node is low (0.36).

**Table 1.** Putative divergence times (TMRCA – time to most recent common ancestor) and 95% highest posterior density (HPD) of the A/Grey seal/England/027661/2017 sequence from the closest related wild bird sequence for each segment, and the posterior probability of the node are shown, along with information about the closest related wild bird sequence including the host, country and year of isolation and subtype of virus. For the MP gene, the closest related sequence according to the maximum-likelihood tree is shown.

| Segment | TMRCA | Lower<br>95% HPD | Upper<br>95% HPD | Posterior<br>probability of<br>node > 0.75 | Host | Country | Year | Subtype |
| --- | --- | --- | --- | --- | --- | --- | --- | --- |
| HA | 2012.74 | 2011.18 | 2014.08 | y | barnacle goose | The Netherlands | 2014 | H3N6 |
| MP | n/a | n/a | n/a | n/a | mallard | The Netherlands | 2010 | H3N2 |
| NA | 1999.53 | 1994.89 | 2003.34 | n | teal/gull | Norway | 2007/2009 | H3N8/H6N8 |
| NP | 2014.98 | 2014.29 | 2015.38 | n | mallard | The Netherlands | 2015 | unknown |
| NS | 2009.56 | 2008.04 | 2010.86 | n | barnacle goose | The Netherlands | 2011 | H6N8 |
| PA | 2013.71 | 2013.1 | 2014.35 | n | mallard | The Netherlands | 2014 | H7N5 |
| PB1 | 2014.28 | 2013.43 | 2015.04 | y | duck | The Netherlands/France | 2015 | H5N1/H5N2/H5N6 |
| PB2 | 2011.45 | 2010.99 | 2011.77 | y | goose/chicken/duck | The Netherlands/France | 2011/2015/2016 | H5N2/H6N2/H7N9/H6N8 |

**Figure 2 (A-G, H).**
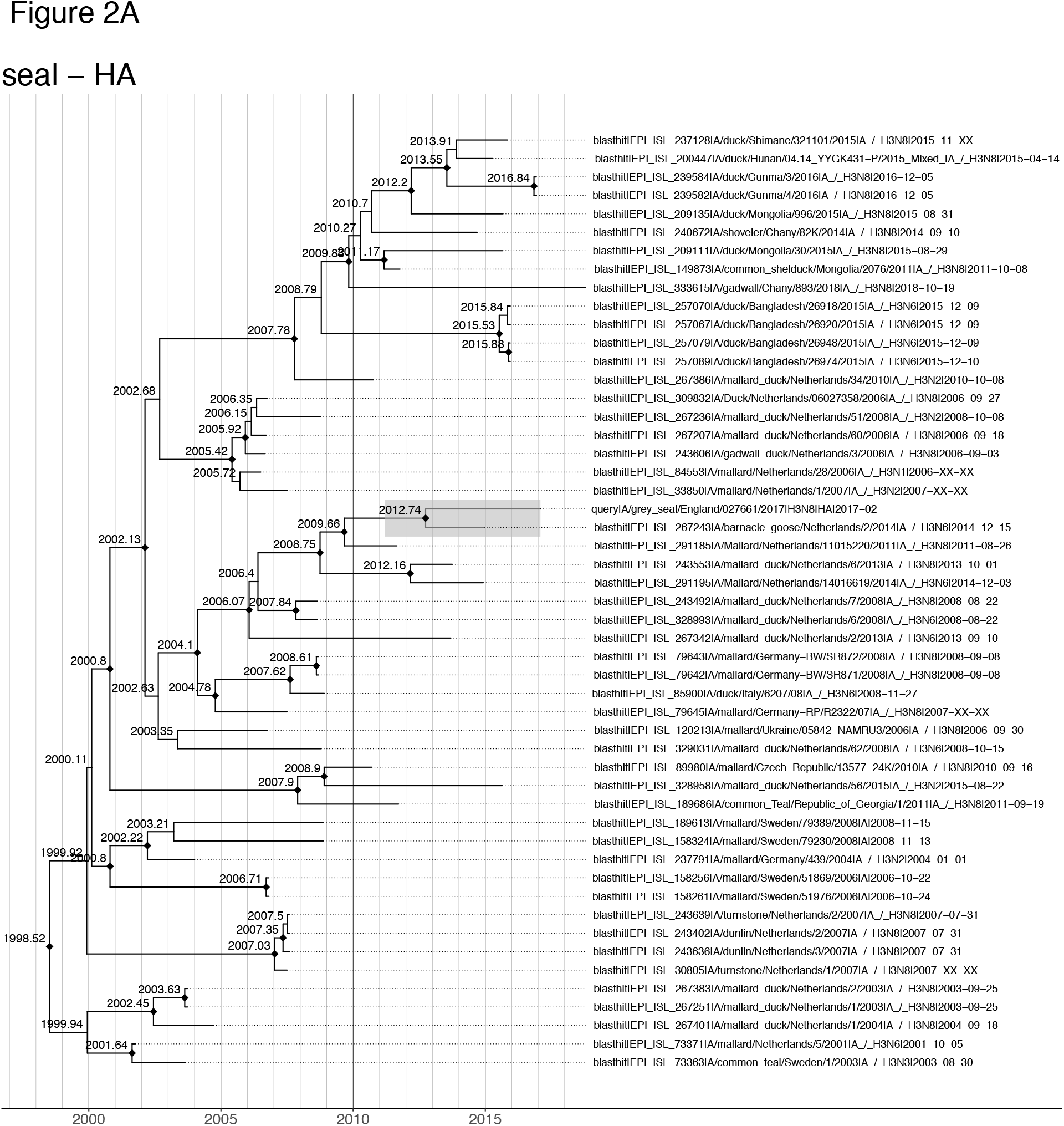
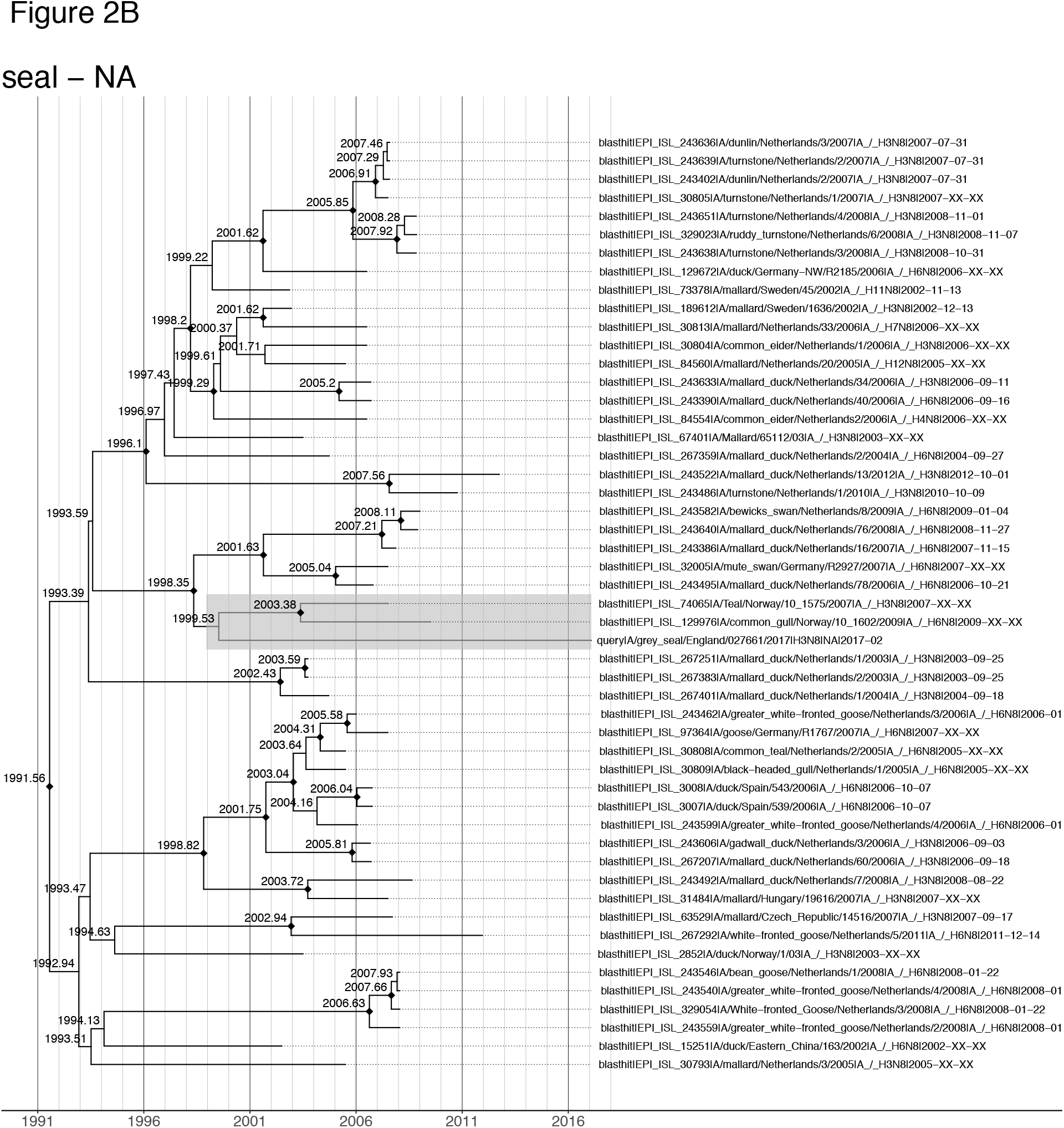
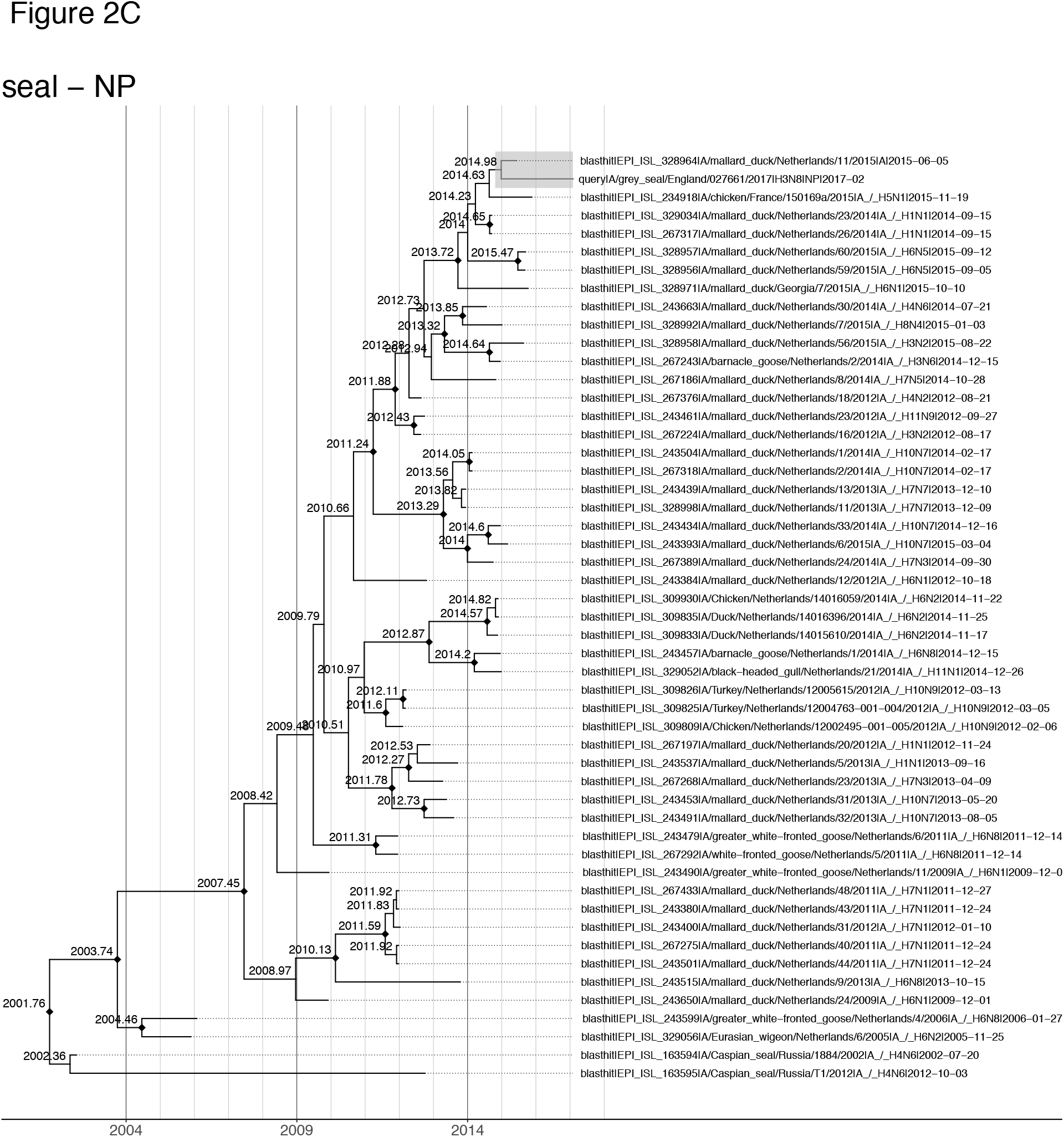
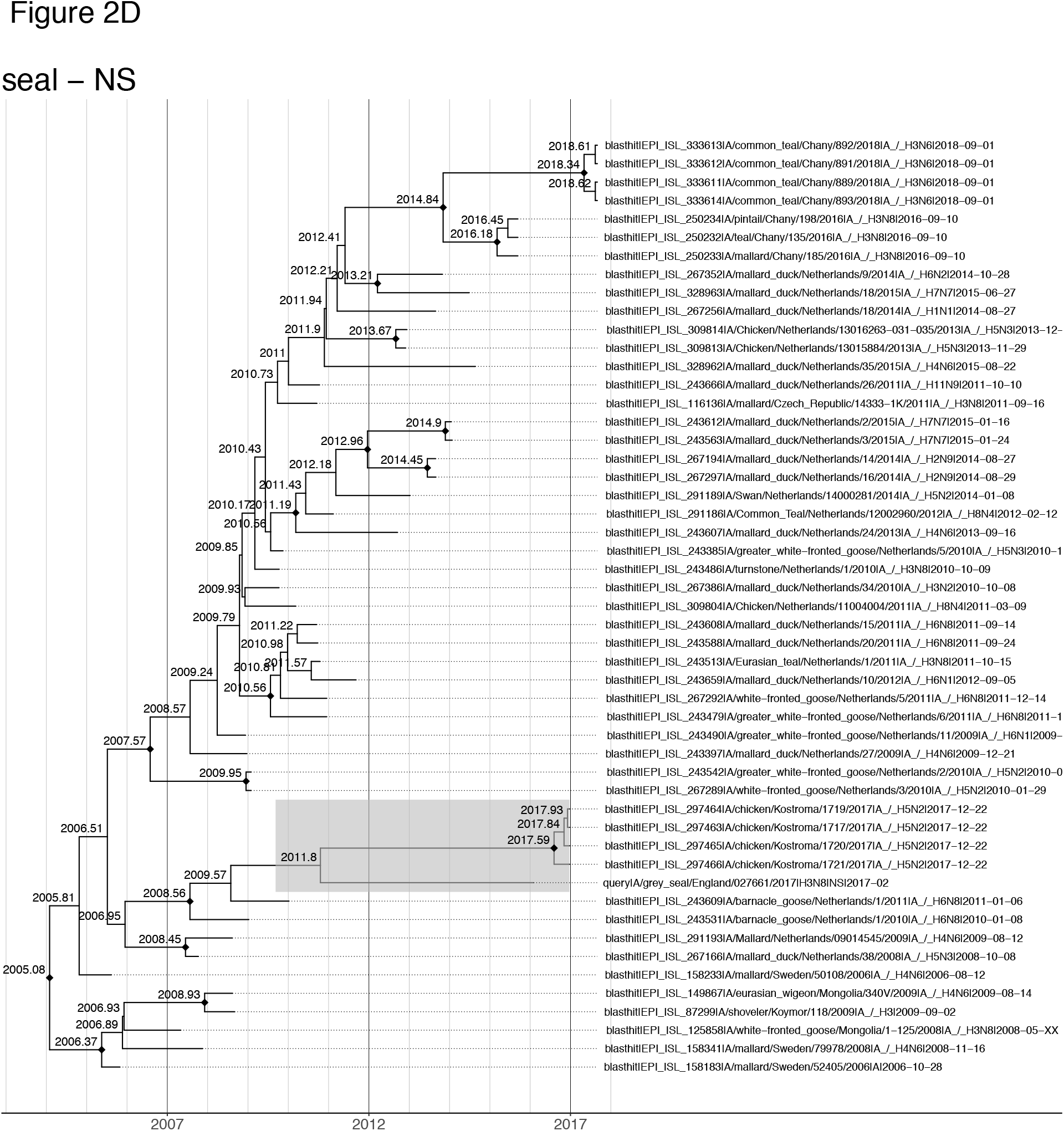
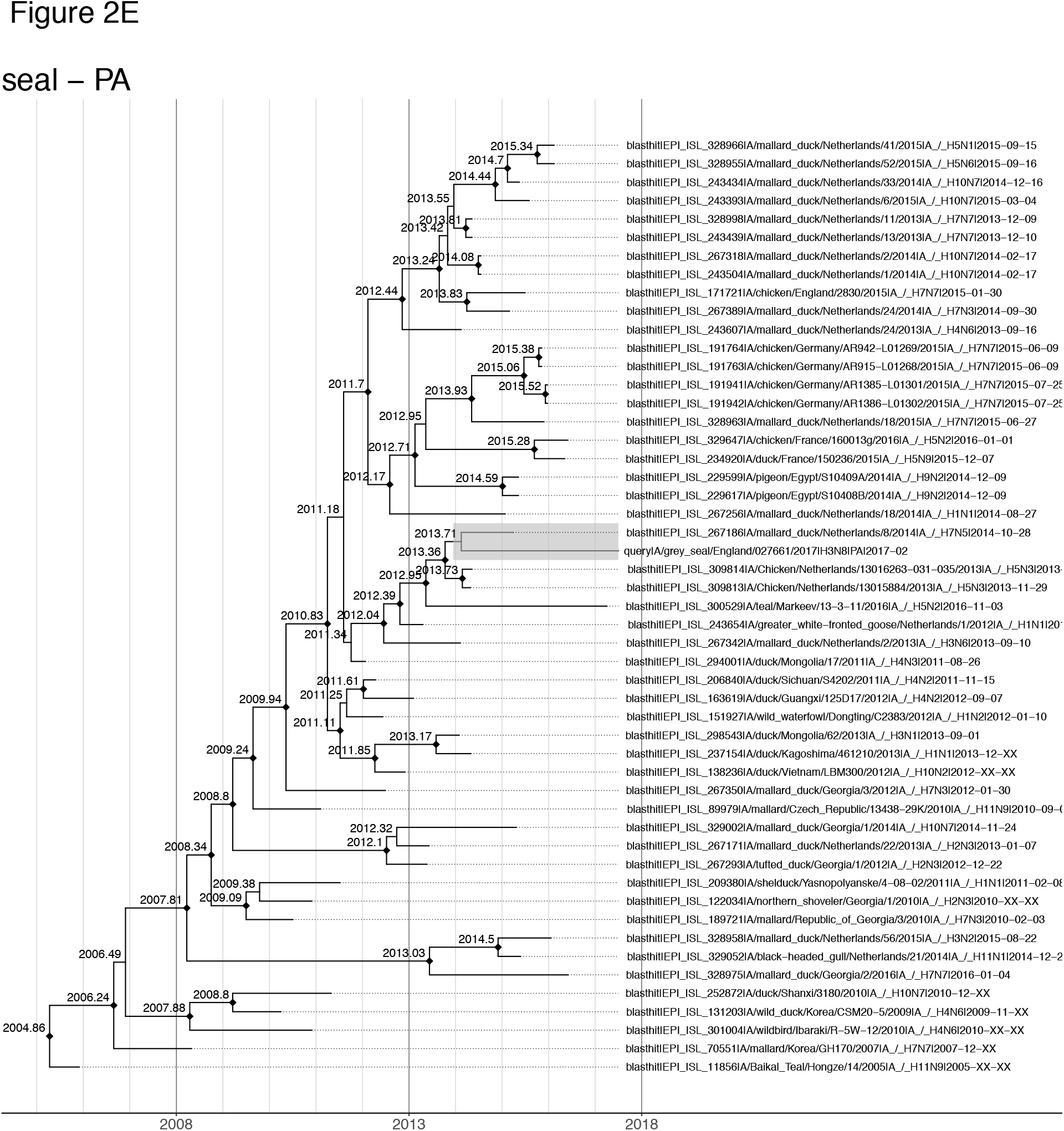
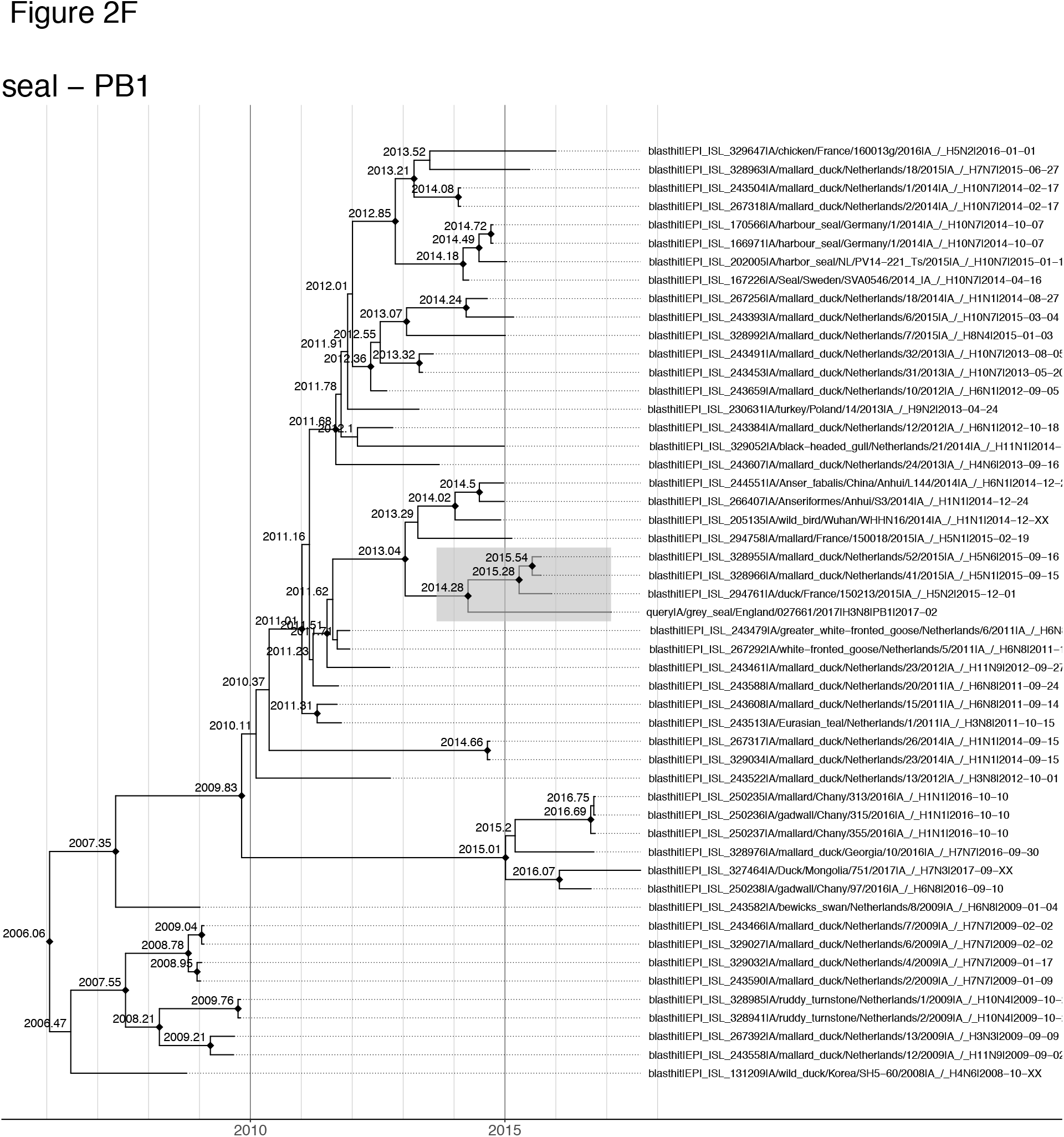
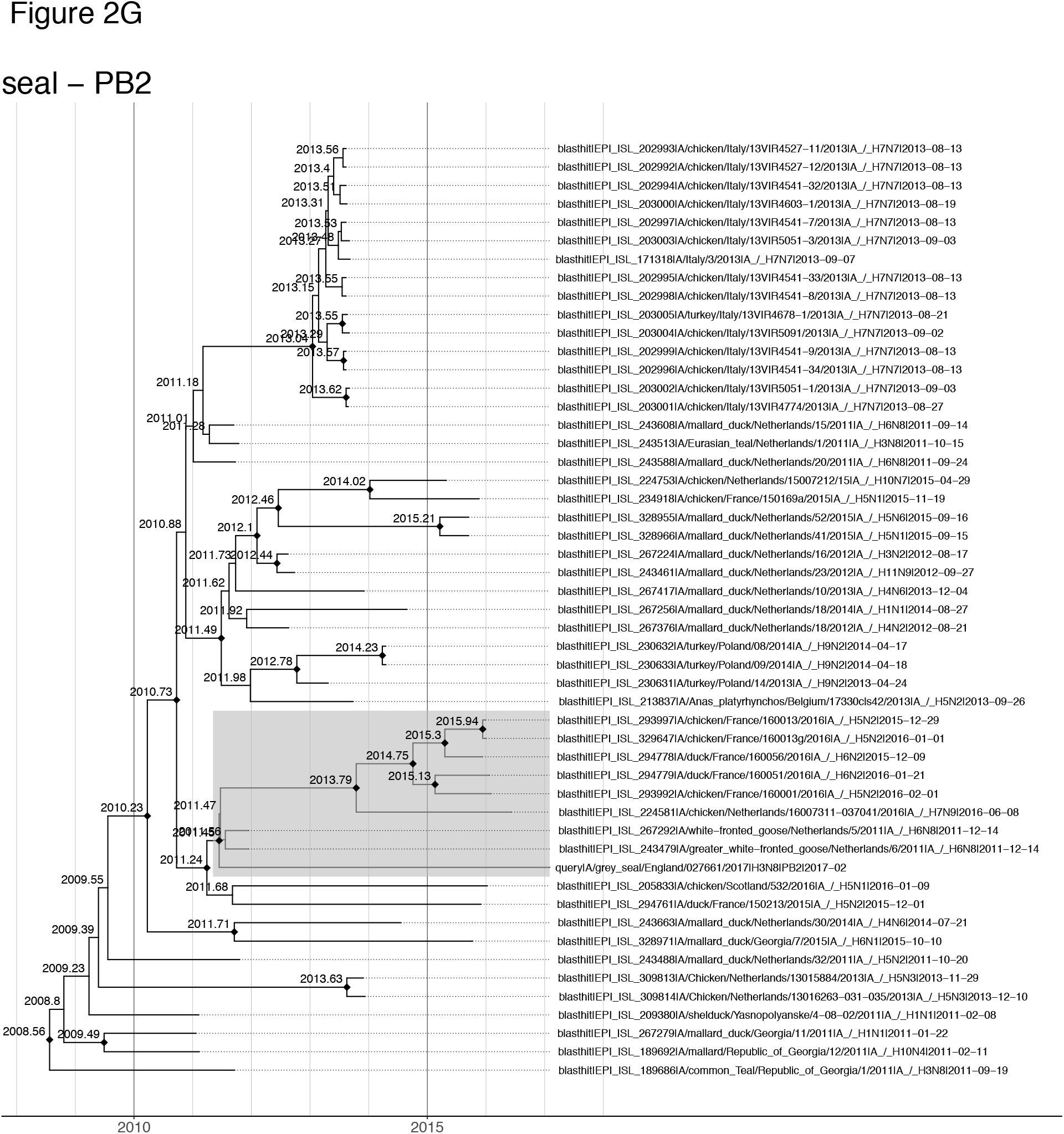
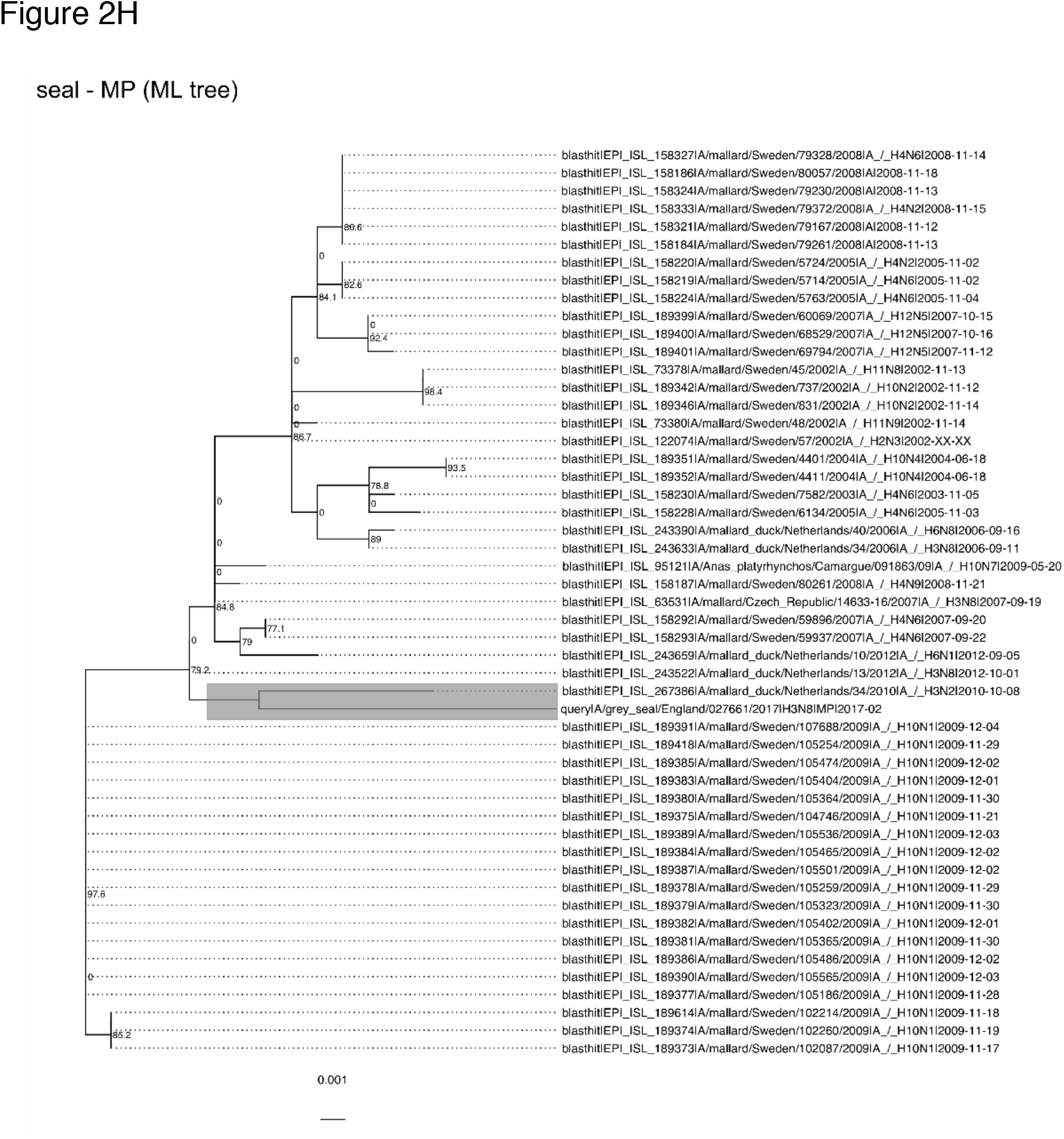
Maximum clade credibility summary trees for BEAST analysis of segment datasets: HA, NA, NP, NS, PA, PB1, PB2 (A-G respectively). Nodes connecting the seal virus tip with the closest related strain(s) are highlighted in grey. Black diamond (◆) shapes at nodes indicate posterior probability > 0.75. Nodes are labelled with node ages as inferred by BEAST. Figure 2H shows the maximum likelihood tree for the MP gene dataset. Support values (approximate likelihood ratio test) are labelled at nodes, and nodes connecting the seal virus tip with the closest related strain is highlighted in grey.

**Figure 3.**
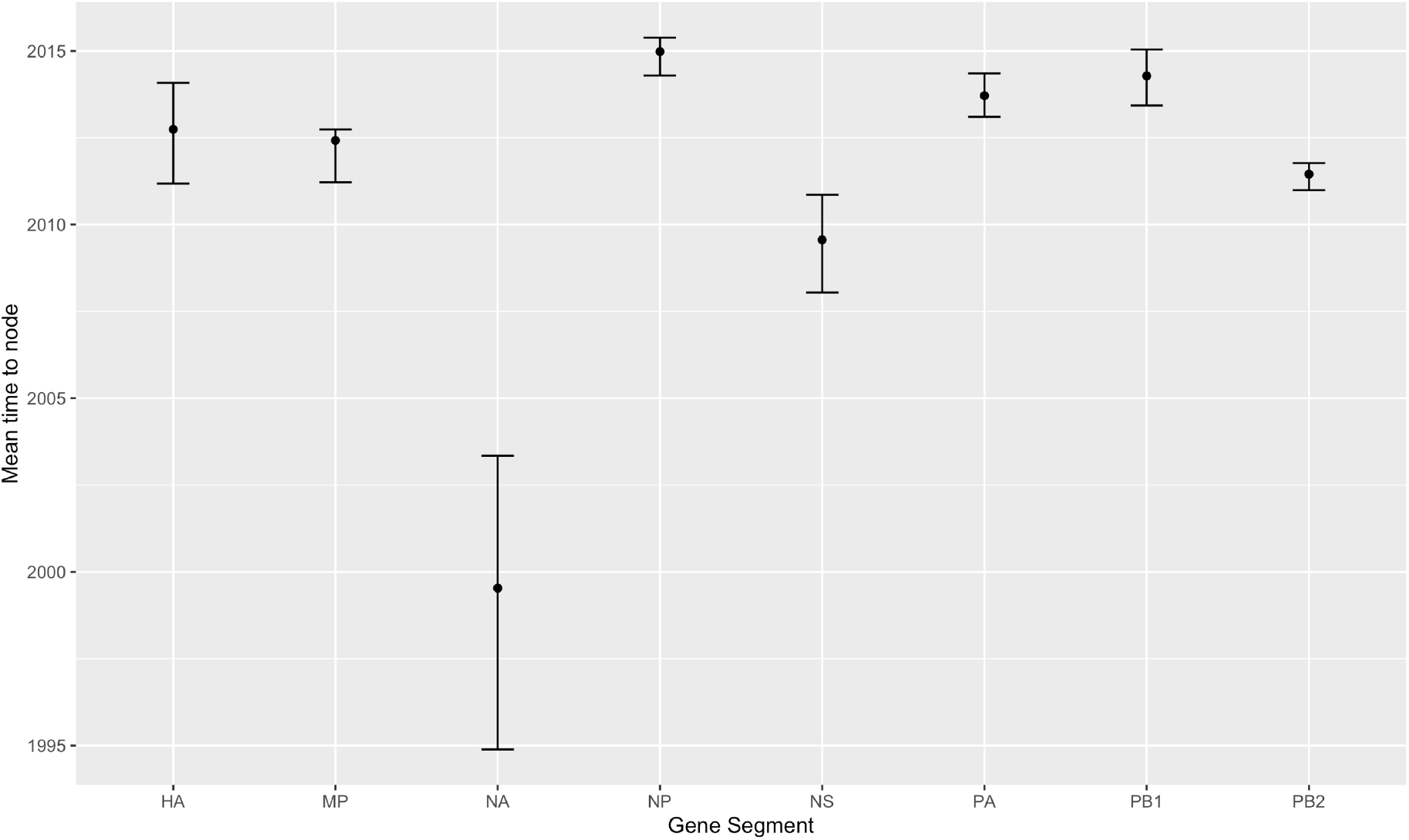
Putative divergence times (TMRCA – time to most recent common ancestor) of the seal sequence from the closest related wild bird sequence for each segment. Error bars indicate 95% highest posterior density.

From the above observations, we conclude that most of the Grey seal virus segments are derived from viruses in wild birds in Northern Europe, possibly via unsampled intermediaries – but not in a single transmission event. In contrast, the 2011 H3N8 Harbour seal virus appears to have been derived from a single source. The closest related strain for all internal gene segments as identified by BLAST and maximum-likelihood trees is the A/American_black_duck/New_Brunswick/03552/2009 (an H4N6 isolated on 2009-09-11). The closest related strains for the glycoproteins are different but this likely indicates a bias for sequencing glycoproteins over the internal genes. They are both H3N8 viruses from the same area (North-eastern US) isolated closer in time to the Harbour seal virus: A/blue-winged_teal/New_Brunswick/00291/2010 isolated 2010-09-14 (for HA) and A/northern_pintail/Minnesota/AI09-4322/2009 isolated 2009-09-12 (for NA). See **Table S2A** and **B** for comparison.

### Substitutions for mammalian adaptation

Previous analyses have indicated that mammalian adaptive mutations can occur in avian viruses when they are transmitted into seals. A study of H10N7 viruses in Harbour seals in Northern Europe, which unlike the present case was demonstrated to have transmitted to other seals and caused an outbreak, showed that mutations were likely to occur early on after transmission to seals and then plateau (Bodewes et al., 2016). We looked for differences between the seal virus and wild bird viruses in the segment amino acid alignments of our datasets, to check if they any had putative adaptive implications.

We found several amino acid substitutions in the seal virus that did not occur in any of the related bird viruses. These substitutions are summarised in Table 2, along with references for those that have been identified in previous studies of mammalian adaptation. Many of these changes occur in the polymerase complex genes (Mänz et al., 2013): D701N in the PB2 segment is a rare mutation, and a hallmark of mammalian adaptation of bird viruses, regardless of genetic background (Liu et al., 2018; Steel et al., 2009)). Liu and Steel et. al. have elucidated that the basis of this adaptation is that it allows for better replication in mammalian cells and showed that it has been associated with increased transmission in ferret experiments. Another mutation in the PB2 gene, at residue 105 in its NP-binding region (Poole et al., 2004) was also found in studies that used phylogenetic modelling (Tamuri et al., 2009) and mutual information statistics (Miotto et al., 2008, 2010). Both the mutations found in the PB1 gene are also of interest. PB1-I517V was found by Tamuri et al. and S678N found in the seal PB1 gene has been associated with increased polymerase activity and virulence in mice (Gabriel et al., 2005).

**Table 2.** Segment-wise list of substitutions found in the seal virus inferred from inspection of alignments. Only residue changes that occur in the seal virus but not in the wild bird viruses are shown. Numbers in square brackets for HA indicate the reference H3-numbering of residues in A/Aichi/2/1968 (Burke and Smith 2014) inferred using the tool at FluDB website: https://tinyurl.com/HAnumbering. For substitutions which have been previously described in a published study, the references are shown in the third column.

| Segment | Amino acid substitutions in seal virus | References |
| --- | --- | --- |
| HA | Y9C [signal peptide] |  |
|  | T64A [48] |  |
|  | I81V [65] |  |
|  | S111G [95] |  |
|  | possible A154T (but might be same RCT --> ACT (T), or GCT (A)) [138] |  |
|  | A/T176S [160] |  |
|  | S235Y [219] |  |
| MP | none |  |
| NA | N/S41D |  |
|  | T383A |  |
| NP | V104M |  |
| NS | none |  |
| PA | V91I |  |
|  | D/G101N |  |
|  | **N/S321I** | (Miotto et al., 2010) |
| PB1 | **I517V** | (Tamuri et al., 2009) |
|  | **S678N** | (Gabriel et al., 2005) |
| PB2 | **T/A105K** | (Tamuri et al., 2009; Miotto et al., 2008) |
|  | D161N |  |
|  | A/S395T |  |
|  | V667I |  |
|  | R/T676I |  |
|  | **D701N** | (Liu et al., 2018; Steel et al., 2009) |
|  | possible R753T (but might be same ASA --> AGA (R) or ACA (T)) |  |

Changes were also found in the HA gene (Table 2), but the implications are less clear. There appear to be no changes in the glycosylation patterns between the HA and NA of the Grey seal virus in comparison to related wild bird viruses (**Table S3A and B**).

We compared all the substitutions with previously described mutations in seal infections, and found that apart from D701N, which was also found in the H3N8 seal virus infection in Massachusetts in 2011 (Anthony et al., 2012), there were no convergent amino acid changes. In the H3 HA gene, we found substitutions in residue 81 (reference H3 numbering - 64), and residue 176 (reference H3 numbering - 160), and these are also changed in the 2011 Massachusetts virus (Anthony et al., 2012) but to different amino acids. The latter mutation was not implicated in receptor binding for the seal viruses as was the case with H5N1 and some human H3N2 viruses, because the glycosylation site is absent in both seal and wild bird viruses (**Table S3A**). The HA of the 2011 Massachusetts virus had an F110S mutation, where the 110 residue has been previously found to be a critical component of the influenza fusion peptide, which may impact replication in mammalian cells (Anthony et al., 2012; Liu et al., 2011). Our reported seal virus retains F at position 110, but whether the mutation in the adjoining residue at S111G (reference H3 numbering 95) has any effect on HA fusion properties is unknown. The presence of Serine at position 66 in the PB1 sequence, which enables production of PB1-F2 (Conenello et al., 2007) was found in the 2011 H3N8 Harbour seal virus but was not seen in this 2017 H3N8 Grey seal virus. Changes at positions 226 and 228 in HA (reference H3 numbering) which can change receptor-binding preferences between avian and mammalian hosts (Connor et al., 1994; Matrosovich et al., 2000), were not found in either of the H3N8 seal viruses, but the H10 equivalent of H3-Q226L was identified in viruses from the 2014-15 H10N7 outbreak in Europe (Dittrich et al., 2018).

## DISCUSSION

In the seal infection case reported here, the animal was referred to a rescue centre because it was stranded and the IAV was detected only incidentally. The vast majority of Grey seals admitted to rehabilitation centres in the UK are pups within the first year of life. Malnutrition is the single most common reason for pinnipeds to be taken into rehabilitation centres (Barnett et al., 2000; Van Bonn, 2015). Wounds are often recorded and hold clinical significance, as in this case, and have been considered predisposing factors for fatal non-specific septicaemia (Baily, 2014). The cause of trauma and wounds may be anthropic (entanglement in fishing nets or gears) or biologic (conspecific aggressiveness or hierarchical to cannibalistic behaviours). Wounds caused by bites of other seals or predators were most often seen (Barnett et al., 2000; Van Bonn, 2015). Pulmonary and nasal parasites are also very well documented clinico-pathological conditions among rescued seal pups, and likely caused the hyperplastic rhinitis identified in this study seal. No histopathological findings consistent with IAV derived damage were detected along the respiratory tract. The immunohistochemistry demonstrated productive viral replication only in nasal mucosa, but not in lower respiratory segments, although the more sensitive RRT-PCR revealed presence of trace amounts of viral RNA within the lower respiratory tract, suggesting that the infection may have been cleared naturally or indicating a passive translocation of antigen from upper respiratory segments. Efficient clearance in healthy Grey seals may explain why previous attempts at sequencing IAV from identified during surveillance were unsuccessful (Puryear et al., 2016).

Puryear et al. discuss the possibility that Grey seals may be more prone to infection due to gain in population density since the marine mammal protection act in the US in 1972, along with more socially gregarious and aggressive behaviour in comparison to Harbour seals, all of which contribute to high pathogen transmissibility. Why we see differences between Grey and Harbour seals in their resistance to diseases caused by viral agents is however unclear. Phylogenetic analyses have been unable to resolve the relationships between different seal species with sufficient support but Grey seals are either placed as a sister group to Caspian seals (*Pusa Caspica*) and separate from Harbour seals, or in a basal position to both *Phoca* and *Pusa* genus (Berta et al., 2018; Fulton and Strobeck, 2006). It might be useful to explore long-term evolutionary history of host-pathogen relationships along with host physiology and immune response differences between different seal species to understand differences in viral pathogenicity.

In our dataset of closest related sequences to the seal virus, we find that while there are several substitutions found in the seal virus which do not occur in the bird viruses; we generally do not find substitutions in any of the bird virus strains that do not occur in other bird viruses too. This likely indicates adaptation to the seal environment, a hypothesis supported by the occurrence of the D701N mutation, a known rare marker of mammalian adaptation (Liu et al., 2018; Steel et al., 2009). D701N is common in canine and horse H3N8 IAV (see **Table S4**), but does not occur in birds, and was found associated with highly-pathogenic H5N1 viruses which infected humans (Gabriel et al., 2005; Jong et al., 2006; Li et al., 2005). We found one reported instance of the D701N mutation in the other avian H3N8 virus that infected pinnipeds, but not in the H10N7 outbreak, despite sustained transmission in seals for several months, nor in any other previously sequenced seal PB2. H3N8 IAV are noted for the ability cross species barriers, so it would be relevant to consider if the subtype of the virus has any influence on the kind of adaptive mutations that occur in the polymerase genes, and if so what sort of mechanisms this might involve. We also note that many of the putative adaptive substitutions occur in the polymerase complex genes, which are increasingly being recognised for their vital role in mediating viral host range (in addition to receptor compatibility with glycoproteins).

The occurrence of adaptive substitutions in the viral sequence could mean that it has been circulating in seals for a certain amount of time during which these substitutions have accumulated. Alternatively, this event might represent an early rapidly-adapting virus from a spill-over event as seen in the H10N7 outbreak (Bodewes et al., 2016). Given the subclinical nature of the infection, we propose the former explanation is more likely to be correct. However, since the case of the seal virus in this study appears to be a singular infection as far as we know, it is difficult to know if and how long the virus has been in seals. Both this study and Bodewes et al. report that despite mammalian adaptation of the virus, continued contact with bird reservoirs allows for further exchange of viral segments between the two hosts. It should be noted that the segment for which this study has tentative evidence of onward transmission (NS) does not appear to have acquired any putative mammalian-adaptive mutations. The nature of the Grey seal reservoir, if it exists, and its relationship with the avian reservoir is currently unknown.

The 2011 North American H3N8 and 2014 European H10N7 viruses which caused outbreaks in Harbour seals were found to have acquired mutations to enable recognition of sialyloligosaccharide receptors found more abundantly in mammalian tissues (SAα2,6Gal) but which retained the ability to interact with avian receptors (SAα2,3Gal). A later detailed structural and functional analysis of the 2011 H3N8 seal HA indicated a true avian receptor binding preference (Yang et al., 2015), as did a mutational analysis of the H10N7 viruses (Dittrich et al., 2018). Although there are some common HA residues that are changed in both the seal virus reported here and the 2011 H3N8 virus, we found limited convergence in the substitutions and residues involved between the different seal viruses, likely attributable to the different genetic backgrounds from different avian sources and possibly differences in host environment of reservoir vs susceptible species. In addition, to the best of our knowledge, ours is the first Grey seal virus sequence that is being made publicly available (via GISAID). It is therefore uncertain whether the type of mutations occurring in a putative reservoir host vis-à-vis avian sequence might be different from those occurring in related hosts with pathogenic outcomes.

Our phylogeny and molecular clock analyses suggest different lineages of source viruses, and time of introduction of different segments (with the caveat of being inferred for a single sequence). It is not surprising to find that the closest sampled and sequenced viruses for each segment are of different subtypes, hosts and times of isolation, given the relatively low surveillance in wild birds, and the extensive reassortment (Lu et al., 2014) i.e., swapping of individual segments into different progeny viruses during propagation after mixed infection with two or more viruses. It would be speculative to propose likely wild bird species donor however. Studies on avian influenza have previously shown that the transmission between birds is directional, e.g., usually in the direction from anseriformes such as ducks into galliformes (chicken) or charadriiformes (gulls) (Venkatesh et al., 2018). In the case of the current infection as well, where we have found unambiguously closest related sequences, they have tended to have been isolated from anseriformes species. The above findings, along with the observation that the closest related viruses were isolated largely from the Netherlands, it is likely that the source of the viral segments come from unsampled locally circulating (in Northern Europe) avian viruses from an anseriformes host. The closest related wild bird viruses to the 2011 H3N8 Harbour seal virus were also isolated from anseriformes species (American black duck). In contrast with the our Grey seal virus however, phylogenetic analysis reveals a highly overlapping set of wild bird strains that are most closely related with Harbour seal virus, an observation that suggests a single spill-over event into a susceptible host.

H3N8 viruses currently circulate in horses and dogs but not humans or pigs. However, different H3 viruses have been found in several species including humans, pigs, horses, dogs, cats, seals, poultry, and wild aquatic birds. They have thus been noted for a particular ability to cross species barrier and cause productive infections. One study that examined the ability of H3N8 viruses from canine, equine, avian, and seal origin to productively infect pigs, demonstrated that avian and seal viruses replicated substantially and caused detectable lesions in inoculated pigs without prior adaptation (Solórzano et al., 2015). It is possible that the ready occurrence of PB2 701N mutation in H3N8 viruses contributes to this ability. We do not have any biological evidence for pre-20th century human influenza, but historical analysis suggests a long association of humans with influenza (Hirsch, 1883; Taubenberger and Morens, 2010), and has uncovered temporal–geographic associations between equine and human influenza-like disease activity documented in Europe in the 16th-18th centuries (Morens and Taubenberger, 2010). Such analysis has also implicated an H3N8 virus in the 1889 pandemic in humans (Morens and Taubenberger, 2010), which makes mammalian-adapted H3N8 viruses of particular interest as IAV pandemic risk candidates.

In this paper we have presented analyses of a case of seal infection with IAV in coastal England, and compared it with previously reported seal IAV infections. This infection provides a small but unique window to understand the ecology of avian-origin viruses that may be circulating and maintained in mammals. Given the mammalian adaptation activity in IAV upon transmission to seals, such infections may be of interest to pandemic surveillance and risk, and help us better understand the determinants of mammalian adaptation in influenza and its complex drivers.

## Notes

#### Summary of Updates

One author was left off on the manuscript pdf - their name has been added. Format of figures and tables changed.

