## Supplemental_legends for "Detection of H3N8 influenza A virus with multiple mammalian-adaptive mutations in a rescued Grey seal (*Halichoerus grypus*) pup"

**Supplementary information legends**

**Figure S1**

Root-to-tip regression for maximum-likelihood (ML) trees generated from each segment (HA, MP, NA, NP, NS, PA, PB1, and PB2) determined using Tempest (v1.5) and

plotted in R (v3.6).

**Table S1**

The number of times a given strain name appears in the blast hits for each seal virus segment, ordered by descending number of segments.

**Table S2 (A-B)**

Table S2A shows the putative closest related wild bird sequence for each segment of the A/harbor_seal/Massachusetts/1/2011 virus, as per maximum likelihood phylogenetic tree generated from the seal virus segment along with the top 50 BLAST hits from GISAID. Columns show node support for the relationship, along with information about the host, location and time of isolation and subtype of virus. The final column shows the other putative closely related strains to the seal virus. Table S2B shows the same information for the A/grey_seal/England/027661/2017 virus.

**Table S3 (A-B)**

Table S3A shows the glycosylation patterns in the grey seal virus (A/grey seal/England/027661/2017) HA compared to the HA of related wild bird viruses (identified by BLAST). For each strain in the first column, the total number of glycosylation sites is shown, followed by (X) for presence and (.) for absence at locations (residue number) in the sequence where Asn-X-Ser or Asn-X-Thr (where X is any amino acid other than Proline) patterns were detected (personal communication from Todd Davis, CDC, USA). Table S3B shows the same information for the NA gene of the seal virus and the closest related NA segments from wild birds.

**Table S4**

The amino acid residue at positions 701 and 627 in the PB2 genes of all non-swine, non-human mammalian viruses found on GISAID (Elbe and Buckland‐Merrett 2017; Shu and McCauley 2017) along with information about the subtype, host and date of isolation of each virus.
