## Supplementary material for "Detection of H3N8 influenza A virus with multiple mammalian-adaptive mutations in a rescued Grey seal (*Halichoerus grypus*) pup": Figure S1

**Figure S1**
Root-to-tip regression for maximum-likelihood (ML) trees generated from each segment (HA, MP, NA, NP, NS, PA, PB1, and PB2) determined using Tempest (v1.5) and plotted in R (v3.6).

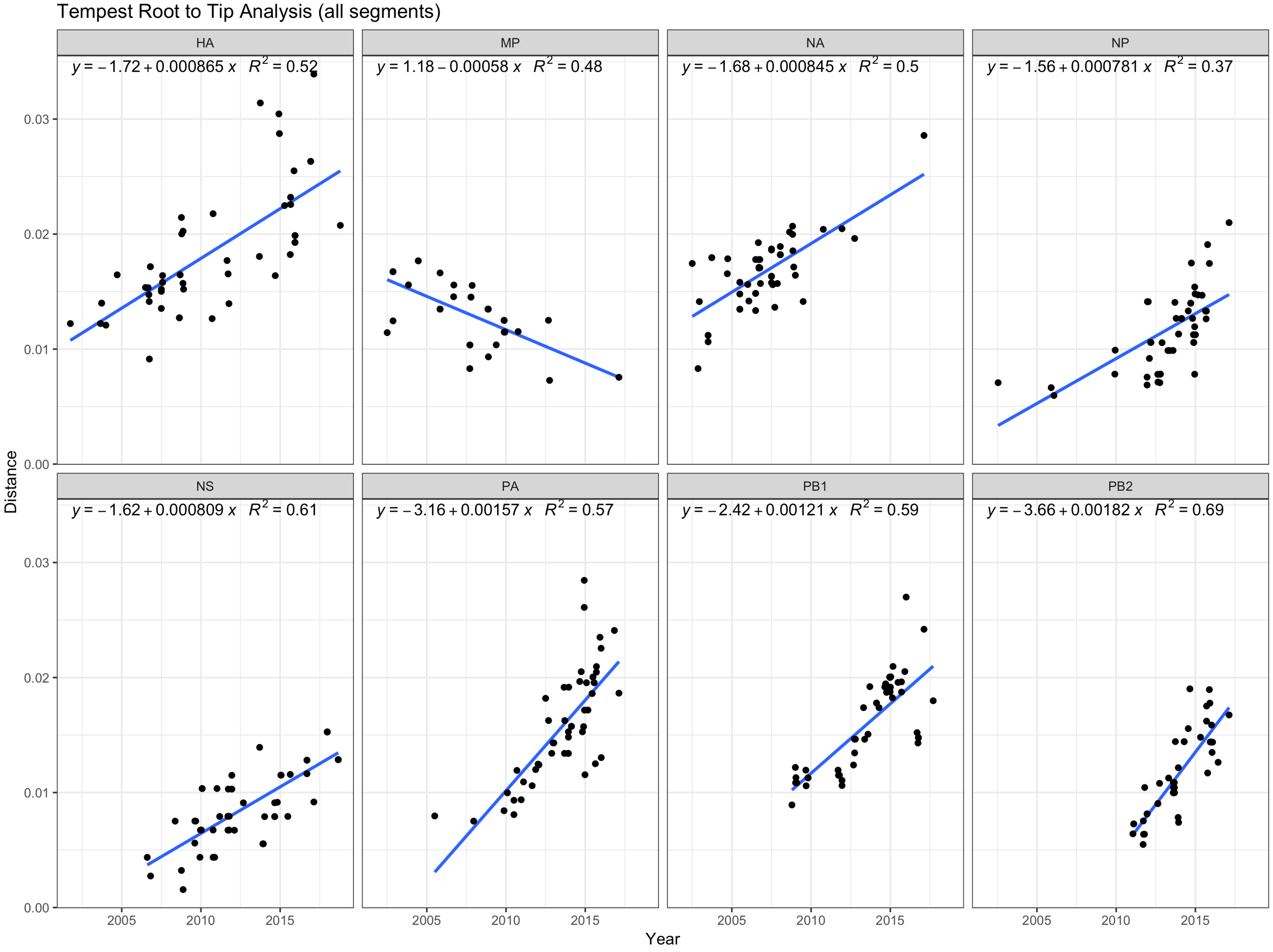
