## Supplementary material for "Detection of H3N8 influenza A virus with multiple mammalian-adaptive mutations in a rescued Grey seal (*Halichoerus grypus*) pup": Table S1

| Isolate ID | Strain Name | Subtype | HA | NA | MP | NP | NS | PA | PB1 | PB2 | Number of Segments in Blast Hits |
|---|---|---|---|---|---|---|---|---|---|---|---|
| EPI_ISL_267292 | A/white-fronted_goose/Netherlands/5/2011 | H6N8 | | present | | present | present | | present | present | 5 |
| EPI_ISL_243479 | A/greater_white-fronted_goose/Netherlands/6/2011 | H6N8 | | | | present | present | | present | present | 4 |
| EPI_ISL_267256 | A/mallard_duck/Netherlands/18/2014 | H1N1 | | | | | present | present | present | present | 4 |
| EPI_ISL_243393 | A/mallard_duck/Netherlands/6/2015 | H10N7 | | | | present | | present | present | | 3 |
| EPI_ISL_243461 | A/mallard_duck/Netherlands/23/2012 | H11N9 | | | | present | | | present | present | 3 |
| EPI_ISL_243504 | A/mallard_duck/Netherlands/1/2014 | H10N7 | | | | present | | present | present | | 3 |
| EPI_ISL_243513 | A/Eurasian_teal/Netherlands/1/2011 | H3N8 | | | | | present | | present | present | 3 |
| EPI_ISL_243522 | A/mallard_duck/Netherlands/13/2012 | H3N8 | | present | present | | | | present | | 3 |
| EPI_ISL_243588 | A/mallard_duck/Netherlands/20/2011 | H6N8 | | | | | present | | present | present | 3 |
| EPI_ISL_243607 | A/mallard_duck/Netherlands/24/2013 | H4N6 | | | | | present | present | present | | 3 |
| EPI_ISL_243608 | A/mallard_duck/Netherlands/15/2011 | H6N8 | | | | | present | | present | present | 3 |
| EPI_ISL_243659 | A/mallard_duck/Netherlands/10/2012 | H6N1 | | | present | | present | | present | | 3 |
| EPI_ISL_267318 | A/mallard_duck/Netherlands/2/2014 | H10N7 | | | | present | | present | present | | 3 |
| EPI_ISL_267386 | A/mallard_duck/Netherlands/34/2010 | H3N2 | present | | present | | present | | | | 3 |
| EPI_ISL_309813 | A/Chicken/Netherlands/13015884/2013 | H5N3 | | | | | present | present | | present | 3 |
| EPI_ISL_309814 | A/Chicken/Netherlands/13016263-031-035/2013 | H5N3 | | | | | present | present | | present | 3 |
| EPI_ISL_328955 | A/mallard_duck/Netherlands/52/2015 | H5N6 | | | | | | present | present | present | 3 |
| EPI_ISL_328958 | A/mallard_duck/Netherlands/56/2015 | H3N2 | present | | | present | | present | | | 3 |
| EPI_ISL_328963 | A/mallard_duck/Netherlands/18/2015 | H7N7 | | | | | present | present | present | | 3 |
| EPI_ISL_328966 | A/mallard_duck/Netherlands/41/2015 | H5N1 | | | | | | present | present | present | 3 |
| EPI_ISL_329052 | A/black-headed_gull/Netherlands/21/2014 | H11N1 | | | | present | | present | present | | 3 |
| EPI_ISL_329647 | A/chicken/France/160013g/2016 | H5N2 | | | | | | present | present | present | 3 |
| EPI_ISL_158324 | A/mallard/Sweden/79230/2008 | unknown | present | | present | | | | | | 2 |
| EPI_ISL_189686 | A/common_Teal/Republic_of_Georgia/1/2011 | H3N8 | present | | | | | | | present | 2 |
| EPI_ISL_209380 | A/shelduck/Yasnopolyanske/4-08-02/2011 | H1N1 | | | | | | present | | present | 2 |
| EPI_ISL_230631 | A/turkey/Poland/14/2013 | H9N2 | | | | | | | present | present | 2 |
| EPI_ISL_234918 | A/chicken/France/150169a/2015 | H5N1 | | | | present | | | | present | 2 |
| EPI_ISL_243384 | A/mallard_duck/Netherlands/12/2012 | H6N1 | | | | present | | | present | | 2 |
| EPI_ISL_243390 | A/mallard_duck/Netherlands/40/2006 | H6N8 | | present | present | | | | | | 2 |
| EPI_ISL_243402 | A/dunlin/Netherlands/2/2007 | H3N8 | present | present | | | | | | | 2 |
| EPI_ISL_243434 | A/mallard_duck/Netherlands/33/2014 | H10N7 | | | | present | | present | | | 2 |
| EPI_ISL_243439 | A/mallard_duck/Netherlands/13/2013 | H7N7 | | | | present | | present | | | 2 |
| EPI_ISL_243453 | A/mallard_duck/Netherlands/31/2013 | H10N7 | | | | present | | | present | | 2 |
| EPI_ISL_243486 | A/turnstone/Netherlands/1/2010 | H3N8 | | present | | | present | | | | 2 |
| EPI_ISL_243490 | A/greater_white-fronted_goose/Netherlands/11/2009 | H6N1 | | | | present | present | | | | 2 |
| EPI_ISL_243491 | A/mallard_duck/Netherlands/32/2013 | H10N7 | | | | present | | | present | | 2 |
| EPI_ISL_243492 | A/mallard_duck/Netherlands/7/2008 | H3N8 | present | present | | | | | | | 2 |
| EPI_ISL_243582 | A/bewicks_swan/Netherlands/8/2009 | H6N8 | | present | | | | | present | | 2 |
| EPI_ISL_243599 | A/greater_white-fronted_goose/Netherlands/4/2006 | H6N8 | | present | | present | | | | | 2 |
| EPI_ISL_243606 | A/gadwall_duck/Netherlands/3/2006 | H3N8 | present | present | | | | | | | 2 |
| EPI_ISL_243633 | A/mallard_duck/Netherlands/34/2006 | H3N8 | | present | present | | | | | | 2 |
| EPI_ISL_243636 | A/dunlin/Netherlands/3/2007 | H3N8 | present | present | | | | | | | 2 |
| EPI_ISL_243639 | A/turnstone/Netherlands/2/2007 | H3N8 | present | present | | | | | | | 2 |
| EPI_ISL_243663 | A/mallard_duck/Netherlands/30/2014 | H4N6 | | | | present | | | | present | 2 |
| EPI_ISL_267186 | A/mallard_duck/Netherlands/8/2014 | H7N5 | | | | present | | present | | | 2 |
| EPI_ISL_267207 | A/mallard_duck/Netherlands/60/2006 | H3N8 | present | present | | | | | | | 2 |
| EPI_ISL_267224 | A/mallard_duck/Netherlands/16/2012 | H3N2 | | | | present | | | | present | 2 |
| EPI_ISL_267243 | A/barnacle_goose/Netherlands/2/2014 | H3N6 | present | | | present | | | | | 2 |
| EPI_ISL_267251 | A/mallard_duck/Netherlands/1/2003 | H3N8 | present | present | | | | | | | 2 |
| EPI_ISL_267317 | A/mallard_duck/Netherlands/26/2014 | H1N1 | | | | present | | | present | | 2 |
| EPI_ISL_267342 | A/mallard_duck/Netherlands/2/2013 | H3N6 | present | | | | | present | | | 2 |
| EPI_ISL_267376 | A/mallard_duck/Netherlands/18/2012 | H4N2 | | | | present | | | | present | 2 |
| EPI_ISL_267383 | A/mallard_duck/Netherlands/2/2003 | H3N8 | present | present | | | | | | | 2 |
| EPI_ISL_267389 | A/mallard_duck/Netherlands/24/2014 | H7N3 | | | | present | | present | | | 2 |
| EPI_ISL_267401 | A/mallard_duck/Netherlands/1/2004 | H3N8 | present | present | | | | | | | 2 |
| EPI_ISL_294761 | A/duck/France/150213/2015 | H5N2 | | | | | | | present | present | 2 |
| EPI_ISL_30805 | A/turnstone/Netherlands/1/2007 | H3N8 | present | present | | | | | | | 2 |
| EPI_ISL_328971 | A/mallard_duck/Georgia/7/2015 | H6N1 | | | | present | | | | present | 2 |
| EPI_ISL_328992 | A/mallard_duck/Netherlands/7/2015 | H8N4 | | | | present | | | present | | 2 |
| EPI_ISL_328998 | A/mallard_duck/Netherlands/11/2013 | H7N7 | | | | present | | present | | | 2 |
| EPI_ISL_329034 | A/mallard_duck/Netherlands/23/2014 | H1N1 | | | | present | | | present | | 2 |
| EPI_ISL_73378 | A/mallard/Sweden/45/2002 | H11N8 | | present | present | | | | | | 2 |
| EPI_ISL_170566 | A/harbour_seal/Germany/1/2014 | H10N7 | | | | | | | present | | 1 |
| EPI_ISL_166971 | A/harbour_seal/Germany/1/2014 | H10N7 | | | | | | | present | | 1 |
| EPI_ISL_116136 | A/mallard/Czech_Republic/14333-1K/2011 | H3N8 | | | | | present | | | | 1 |
| EPI_ISL_11856 | A/Baikal_Teal/Hongze/14/2005 | H11N9 | | | | | | present | | | 1 |
| EPI_ISL_120213 | A/mallard/Ukraine/05842-NAMRU3/2006 | H3N8 | present | | | | | | | | 1 |
| EPI_ISL_122034 | A/northern_shoveler/Georgia/1/2010 | H2N3 | | | | | | present | | | 1 |
| EPI_ISL_122074 | A/mallard/Sweden/57/2002 | H2N3 | | | present | | | | | | 1 |
| EPI_ISL_125858 | A/white-fronted_goose/Mongolia/1-125/2008 | H3N8 | | | | | present | | | | 1 |
| EPI_ISL_129672 | A/duck/Germany-NW/R2185/2006 | H6N8 | | present | | | | | | | 1 |
| EPI_ISL_129976 | A/common_gull/Norway/10_1602/2009 | H6N8 | | present | | | | | | | 1 |
| EPI_ISL_131203 | A/wild_duck/Korea/CSM20-5/2009 | H4N6 | | | | | | present | | | 1 |
| EPI_ISL_131209 | A/wild_duck/Korea/SH5-60/2008 | H4N6 | | | | | | | present | | 1 |
| EPI_ISL_138236 | A/duck/Vietnam/LBM300/2012 | H10N2 | | | | | | present | | | 1 |
| EPI_ISL_149867 | A/eurasian_wigeon/Mongolia/340V/2009 | H4N6 | | | | | present | | | | 1 |
| EPI_ISL_149873 | A/common_shelduck/Mongolia/2076/2011 | H3N8 | present | | | | | | | | 1 |
| EPI_ISL_151927 | A/wild_waterfowl/Dongting/C2383/2012 | H1N2 | | | | | | present | | | 1 |
| EPI_ISL_15251 | A/duck/Eastern_China/163/2002 | H6N8 | | present | | | | | | | 1 |
| EPI_ISL_158183 | A/mallard/Sweden/52405/2006 | unknown | | | | | present | | | | 1 |
| EPI_ISL_158184 | A/mallard/Sweden/79261/2008 | unknown | | | present | | | | | | 1 |
| EPI_ISL_158186 | A/mallard/Sweden/80057/2008 | unknown | | | present | | | | | | 1 |
| EPI_ISL_158187 | A/mallard/Sweden/80261/2008 | H4N9 | | | present | | | | | | 1 |
| EPI_ISL_158219 | A/mallard/Sweden/5714/2005 | H4N6 | | | present | | | | | | 1 |
| EPI_ISL_158220 | A/mallard/Sweden/5724/2005 | H4N2 | | | present | | | | | | 1 |
| EPI_ISL_158224 | A/mallard/Sweden/5763/2005 | H4N6 | | | present | | | | | | 1 |
| EPI_ISL_158228 | A/mallard/Sweden/6134/2005 | H4N6 | | | present | | | | | | 1 |
| EPI_ISL_158230 | A/mallard/Sweden/7582/2003 | H4N6 | | | present | | | | | | 1 |
| EPI_ISL_158233 | A/mallard/Sweden/50108/2006 | H4N6 | | | | | present | | | | 1 |
| EPI_ISL_158256 | A/mallard/Sweden/51869/2006 | unknown | present | | | | | | | | 1 |
| EPI_ISL_158261 | A/mallard/Sweden/51976/2006 | unknown | present | | | | | | | | 1 |
| EPI_ISL_158292 | A/mallard/Sweden/59896/2007 | H4N6 | | | present | | | | | | 1 |
| EPI_ISL_158293 | A/mallard/Sweden/59937/2007 | H4N6 | | | present | | | | | | 1 |
| EPI_ISL_158321 | A/mallard/Sweden/79167/2008 | unknown | | | present | | | | | | 1 |
| EPI_ISL_158327 | A/mallard/Sweden/79328/2008 | H4N6 | | | present | | | | | | 1 |
| EPI_ISL_158333 | A/mallard/Sweden/79372/2008 | H4N2 | | | present | | | | | | 1 |
| EPI_ISL_158341 | A/mallard/Sweden/79978/2008 | H4N6 | | | | | present | | | | 1 |
| EPI_ISL_163594 | A/Caspian_seal/Russia/1884/2002 | H4N6 | | | | present | | | | | 1 |
| EPI_ISL_163595 | A/Caspian_seal/Russia/T1/2012 | H4N6 | | | | present | | | | | 1 |
| EPI_ISL_163619 | A/duck/Guangxi/125D17/2012 | H4N2 | | | | | | present | | | 1 |
| EPI_ISL_167226 | A/Seal/Sweden/SVA0546/2014_ | H10N7 | | | | | | | present | | 1 |
| EPI_ISL_171318 | A/Italy/3/2013 | H7N7 | | | | | | | | present | 1 |
| EPI_ISL_171721 | A/chicken/England/2830/2015 | H7N7 | | | | | | present | | | 1 |
| EPI_ISL_189342 | A/mallard/Sweden/737/2002 | H10N2 | | | present | | | | | | 1 |
| EPI_ISL_189346 | A/mallard/Sweden/831/2002 | H10N2 | | | present | | | | | | 1 |
| EPI_ISL_189351 | A/mallard/Sweden/4401/2004 | H10N4 | | | present | | | | | | 1 |
| EPI_ISL_189352 | A/mallard/Sweden/4411/2004 | H10N4 | | | present | | | | | | 1 |
| EPI_ISL_189373 | A/mallard/Sweden/102087/2009 | H10N1 | | | present | | | | | | 1 |
| EPI_ISL_189374 | A/mallard/Sweden/102260/2009 | H10N1 | | | present | | | | | | 1 |
| EPI_ISL_189375 | A/mallard/Sweden/104746/2009 | H10N1 | | | present | | | | | | 1 |
| EPI_ISL_189377 | A/mallard/Sweden/105186/2009 | H10N1 | | | present | | | | | | 1 |
| EPI_ISL_189378 | A/mallard/Sweden/105259/2009 | H10N1 | | | present | | | | | | 1 |
| EPI_ISL_189379 | A/mallard/Sweden/105323/2009 | H10N1 | | | present | | | | | | 1 |
| EPI_ISL_189380 | A/mallard/Sweden/105364/2009 | H10N1 | | | present | | | | | | 1 |
| EPI_ISL_189381 | A/mallard/Sweden/105365/2009 | H10N1 | | | present | | | | | | 1 |
| EPI_ISL_189382 | A/mallard/Sweden/105402/2009 | H10N1 | | | present | | | | | | 1 |
| EPI_ISL_189383 | A/mallard/Sweden/105404/2009 | H10N1 | | | present | | | | | | 1 |
| EPI_ISL_189384 | A/mallard/Sweden/105465/2009 | H10N1 | | | present | | | | | | 1 |
| EPI_ISL_189385 | A/mallard/Sweden/105474/2009 | H10N1 | | | present | | | | | | 1 |
| EPI_ISL_189386 | A/mallard/Sweden/105486/2009 | H10N1 | | | present | | | | | | 1 |
| EPI_ISL_189387 | A/mallard/Sweden/105501/2009 | H10N1 | | | present | | | | | | 1 |
| EPI_ISL_189389 | A/mallard/Sweden/105536/2009 | H10N1 | | | present | | | | | | 1 |
| EPI_ISL_189390 | A/mallard/Sweden/105565/2009 | H10N1 | | | present | | | | | | 1 |
| EPI_ISL_189391 | A/mallard/Sweden/107688/2009 | H10N1 | | | present | | | | | | 1 |
| EPI_ISL_189399 | A/mallard/Sweden/60069/2007 | H12N5 | | | present | | | | | | 1 |
| EPI_ISL_189400 | A/mallard/Sweden/68529/2007 | H12N5 | | | present | | | | | | 1 |
| EPI_ISL_189401 | A/mallard/Sweden/69794/2007 | H12N5 | | | present | | | | | | 1 |
| EPI_ISL_189418 | A/mallard/Sweden/105254/2009 | H10N1 | | | present | | | | | | 1 |
| EPI_ISL_189612 | A/mallard/Sweden/1636/2002 | H3N8 | | present | | | | | | | 1 |
| EPI_ISL_189613 | A/mallard/Sweden/79389/2008 | unknown | present | | | | | | | | 1 |
| EPI_ISL_189614 | A/mallard/Sweden/102214/2009 | H10N1 | | | present | | | | | | 1 |
| EPI_ISL_189692 | A/mallard/Republic_of_Georgia/12/2011 | H10N4 | | | | | | | | present | 1 |
| EPI_ISL_189721 | A/mallard/Republic_of_Georgia/3/2010 | H7N3 | | | | | | present | | | 1 |
| EPI_ISL_191763 | A/chicken/Germany/AR915-L01268/2015 | H7N7 | | | | | | present | | | 1 |
| EPI_ISL_191764 | A/chicken/Germany/AR942-L01269/2015 | H7N7 | | | | | | present | | | 1 |
| EPI_ISL_191941 | A/chicken/Germany/AR1385-L01301/2015 | H7N7 | | | | | | present | | | 1 |
| EPI_ISL_191942 | A/chicken/Germany/AR1386-L01302/2015 | H7N7 | | | | | | present | | | 1 |
| EPI_ISL_200447 | A/duck/Hunan/04.14_YYGK431-P/2015_Mixed_ | H3N8 | present | | | | | | | | 1 |
| EPI_ISL_202005 | A/harbor_seal/NL/PV14-221_Ts/2015 | H10N7 | | | | | | | present | | 1 |
| EPI_ISL_202992 | A/chicken/Italy/13VIR4527-12/2013 | H7N7 | | | | | | | | present | 1 |
| EPI_ISL_202993 | A/chicken/Italy/13VIR4527-11/2013 | H7N7 | | | | | | | | present | 1 |
| EPI_ISL_202994 | A/chicken/Italy/13VIR4541-32/2013 | H7N7 | | | | | | | | present | 1 |
| EPI_ISL_202995 | A/chicken/Italy/13VIR4541-33/2013 | H7N7 | | | | | | | | present | 1 |
| EPI_ISL_202996 | A/chicken/Italy/13VIR4541-34/2013 | H7N7 | | | | | | | | present | 1 |
| EPI_ISL_202997 | A/chicken/Italy/13VIR4541-7/2013 | H7N7 | | | | | | | | present | 1 |
| EPI_ISL_202998 | A/chicken/Italy/13VIR4541-8/2013 | H7N7 | | | | | | | | present | 1 |
| EPI_ISL_202999 | A/chicken/Italy/13VIR4541-9/2013 | H7N7 | | | | | | | | present | 1 |
| EPI_ISL_203000 | A/chicken/Italy/13VIR4603-1/2013 | H7N7 | | | | | | | | present | 1 |
| EPI_ISL_203001 | A/chicken/Italy/13VIR4774/2013 | H7N7 | | | | | | | | present | 1 |
| EPI_ISL_203002 | A/chicken/Italy/13VIR5051-1/2013 | H7N7 | | | | | | | | present | 1 |
| EPI_ISL_203003 | A/chicken/Italy/13VIR5051-3/2013 | H7N7 | | | | | | | | present | 1 |
| EPI_ISL_203004 | A/chicken/Italy/13VIR5091/2013 | H7N7 | | | | | | | | present | 1 |
| EPI_ISL_203005 | A/turkey/Italy/13VIR4678-1/2013 | H7N7 | | | | | | | | present | 1 |
| EPI_ISL_205135 | A/wild_bird/Wuhan/WHHN16/2014 | H1N1 | | | | | | | present | | 1 |
| EPI_ISL_205833 | A/chicken/Scotland/532/2016 | H5N1 | | | | | | | | present | 1 |
| EPI_ISL_206840 | A/duck/Sichuan/S4202/2011 | H4N2 | | | | | | present | | | 1 |
| EPI_ISL_209111 | A/duck/Mongolia/30/2015 | H3N8 | present | | | | | | | | 1 |
| EPI_ISL_209135 | A/duck/Mongolia/996/2015 | H3N8 | present | | | | | | | | 1 |
| EPI_ISL_213837 | A/Anas_platyrhynchos/Belgium/17330cls42/2013 | H5N2 | | | | | | | | present | 1 |
| EPI_ISL_224581 | A/chicken/Netherlands/16007311-037041/2016 | H7N9 | | | | | | | | present | 1 |
| EPI_ISL_224753 | A/chicken/Netherlands/15007212/15 | H10N7 | | | | | | | | present | 1 |
| EPI_ISL_229599 | A/pigeon/Egypt/S10409A/2014 | H9N2 | | | | | | present | | | 1 |
| EPI_ISL_229617 | A/pigeon/Egypt/S10408B/2014 | H9N2 | | | | | | present | | | 1 |
| EPI_ISL_230632 | A/turkey/Poland/08/2014 | H9N2 | | | | | | | | present | 1 |
| EPI_ISL_230633 | A/turkey/Poland/09/2014 | H9N2 | | | | | | | | present | 1 |
| EPI_ISL_234920 | A/duck/France/150236/2015 | H5N9 | | | | | | present | | | 1 |
| EPI_ISL_237128 | A/duck/Shimane/321101/2015 | H3N8 | present | | | | | | | | 1 |
| EPI_ISL_237154 | A/duck/Kagoshima/461210/2013 | H1N1 | | | | | | present | | | 1 |
| EPI_ISL_237791 | A/mallard/Germany/439/2004 | H3N2 | present | | | | | | | | 1 |
| EPI_ISL_239582 | A/duck/Gunma/4/2016 | H3N8 | present | | | | | | | | 1 |
| EPI_ISL_239584 | A/duck/Gunma/3/2016 | H3N8 | present | | | | | | | | 1 |
| EPI_ISL_240672 | A/shoveler/Chany/82K/2014 | H3N8 | present | | | | | | | | 1 |
| EPI_ISL_243380 | A/mallard_duck/Netherlands/43/2011 | H7N1 | | | | present | | | | | 1 |
| EPI_ISL_243385 | A/greater_white-fronted_goose/Netherlands/5/2010 | H5N3 | | | | | present | | | | 1 |
| EPI_ISL_243386 | A/mallard_duck/Netherlands/16/2007 | H6N8 | | present | | | | | | | 1 |
| EPI_ISL_243397 | A/mallard_duck/Netherlands/27/2009 | H4N6 | | | | | present | | | | 1 |
| EPI_ISL_243400 | A/mallard_duck/Netherlands/31/2012 | H7N1 | | | | present | | | | | 1 |
| EPI_ISL_243457 | A/barnacle_goose/Netherlands/1/2014 | H6N8 | | | | present | | | | | 1 |
| EPI_ISL_243462 | A/greater_white-fronted_goose/Netherlands/3/2006 | H6N8 | | present | | | | | | | 1 |
| EPI_ISL_243466 | A/mallard_duck/Netherlands/7/2009 | H7N7 | | | | | | | present | | 1 |
| EPI_ISL_243488 | A/mallard_duck/Netherlands/32/2011 | H5N2 | | | | | | | | present | 1 |
| EPI_ISL_243495 | A/mallard_duck/Netherlands/78/2006 | H6N8 | | present | | | | | | | 1 |
| EPI_ISL_243501 | A/mallard_duck/Netherlands/44/2011 | H7N1 | | | | present | | | | | 1 |
| EPI_ISL_243515 | A/mallard_duck/Netherlands/9/2013 | H6N8 | | | | present | | | | | 1 |
| EPI_ISL_243531 | A/barnacle_goose/Netherlands/1/2010 | H6N8 | | | | | present | | | | 1 |
| EPI_ISL_243537 | A/mallard_duck/Netherlands/5/2013 | H1N1 | | | | present | | | | | 1 |
| EPI_ISL_243540 | A/greater_white-fronted_goose/Netherlands/4/2008 | H6N8 | | present | | | | | | | 1 |
| EPI_ISL_243542 | A/greater_white-fronted_goose/Netherlands/2/2010 | H5N2 | | | | | present | | | | 1 |
| EPI_ISL_243546 | A/bean_goose/Netherlands/1/2008 | H6N8 | | present | | | | | | | 1 |
| EPI_ISL_243553 | A/mallard_duck/Netherlands/6/2013 | H3N8 | present | | | | | | | | 1 |
| EPI_ISL_243558 | A/mallard_duck/Netherlands/12/2009 | H11N9 | | | | | | | present | | 1 |
| EPI_ISL_243559 | A/greater_white-fronted_goose/Netherlands/2/2008 | H6N8 | | present | | | | | | | 1 |
| EPI_ISL_243563 | A/mallard_duck/Netherlands/3/2015 | H7N7 | | | | | present | | | | 1 |
| EPI_ISL_243590 | A/mallard_duck/Netherlands/2/2009 | H7N7 | | | | | | | present | | 1 |
| EPI_ISL_243609 | A/barnacle_goose/Netherlands/1/2011 | H6N8 | | | | | present | | | | 1 |
| EPI_ISL_243612 | A/mallard_duck/Netherlands/2/2015 | H7N7 | | | | | present | | | | 1 |
| EPI_ISL_243638 | A/turnstone/Netherlands/3/2008 | H3N8 | | present | | | | | | | 1 |
| EPI_ISL_243640 | A/mallard_duck/Netherlands/76/2008 | H6N8 | | present | | | | | | | 1 |
| EPI_ISL_243650 | A/mallard_duck/Netherlands/24/2009 | H6N1 | | | | present | | | | | 1 |
| EPI_ISL_243651 | A/turnstone/Netherlands/4/2008 | H3N8 | | present | | | | | | | 1 |
| EPI_ISL_243654 | A/greater_white-fronted_goose/Netherlands/1/2012 | H1N1 | | | | | | present | | | 1 |
| EPI_ISL_243666 | A/mallard_duck/Netherlands/26/2011 | H11N9 | | | | | present | | | | 1 |
| EPI_ISL_244551 | A/Anser_fabalis/China/Anhui/L144/2014 | H6N1 | | | | | | | present | | 1 |
| EPI_ISL_250232 | A/teal/Chany/135/2016 | H3N8 | | | | | present | | | | 1 |
| EPI_ISL_250233 | A/mallard/Chany/185/2016 | H3N8 | | | | | present | | | | 1 |
| EPI_ISL_250234 | A/pintail/Chany/198/2016 | H3N8 | | | | | present | | | | 1 |
| EPI_ISL_250235 | A/mallard/Chany/313/2016 | H1N1 | | | | | | | present | | 1 |
| EPI_ISL_250236 | A/gadwall/Chany/315/2016 | H1N1 | | | | | | | present | | 1 |
| EPI_ISL_250237 | A/mallard/Chany/355/2016 | H1N1 | | | | | | | present | | 1 |
| EPI_ISL_250238 | A/gadwall/Chany/97/2016 | H6N8 | | | | | | | present | | 1 |
| EPI_ISL_252872 | A/duck/Shanxi/3180/2010 | H10N7 | | | | | | present | | | 1 |
| EPI_ISL_257067 | A/duck/Bangladesh/26920/2015 | H3N6 | present | | | | | | | | 1 |
| EPI_ISL_257070 | A/duck/Bangladesh/26918/2015 | H3N6 | present | | | | | | | | 1 |
| EPI_ISL_257079 | A/duck/Bangladesh/26948/2015 | H3N6 | present | | | | | | | | 1 |
| EPI_ISL_257089 | A/duck/Bangladesh/26974/2015 | H3N6 | present | | | | | | | | 1 |
| EPI_ISL_266407 | A/Anseriformes/Anhui/S3/2014 | H1N1 | | | | | | | present | | 1 |
| EPI_ISL_267166 | A/mallard_duck/Netherlands/38/2008 | H5N3 | | | | | present | | | | 1 |
| EPI_ISL_267171 | A/mallard_duck/Netherlands/22/2013 | H2N3 | | | | | | present | | | 1 |
| EPI_ISL_267194 | A/mallard_duck/Netherlands/14/2014 | H2N9 | | | | | present | | | | 1 |
| EPI_ISL_267197 | A/mallard_duck/Netherlands/20/2012 | H1N1 | | | | present | | | | | 1 |
| EPI_ISL_267236 | A/mallard_duck/Netherlands/51/2008 | H3N2 | present | | | | | | | | 1 |
| EPI_ISL_267268 | A/mallard_duck/Netherlands/23/2013 | H7N3 | | | | present | | | | | 1 |
| EPI_ISL_267275 | A/mallard_duck/Netherlands/40/2011 | H7N1 | | | | present | | | | | 1 |
| EPI_ISL_267279 | A/mallard_duck/Georgia/11/2011 | H1N1 | | | | | | | | present | 1 |
| EPI_ISL_267289 | A/white-fronted_goose/Netherlands/3/2010 | H5N2 | | | | | present | | | | 1 |
| EPI_ISL_267293 | A/tufted_duck/Georgia/1/2012 | H2N3 | | | | | | present | | | 1 |
| EPI_ISL_267297 | A/mallard_duck/Netherlands/16/2014 | H2N9 | | | | | present | | | | 1 |
| EPI_ISL_267350 | A/mallard_duck/Georgia/3/2012 | H7N3 | | | | | | present | | | 1 |
| EPI_ISL_267352 | A/mallard_duck/Netherlands/9/2014 | H6N2 | | | | | present | | | | 1 |
| EPI_ISL_267359 | A/mallard_duck/Netherlands/2/2004 | H6N8 | | present | | | | | | | 1 |
| EPI_ISL_267392 | A/mallard_duck/Netherlands/13/2009 | H3N3 | | | | | | | present | | 1 |
| EPI_ISL_267417 | A/mallard_duck/Netherlands/10/2013 | H4N6 | | | | | | | | present | 1 |
| EPI_ISL_267433 | A/mallard_duck/Netherlands/48/2011 | H7N1 | | | | present | | | | | 1 |
| EPI_ISL_2852 | A/duck/Norway/1/03 | H3N8 | | present | | | | | | | 1 |
| EPI_ISL_291185 | A/Mallard/Netherlands/11015220/2011 | H3N8 | present | | | | | | | | 1 |
| EPI_ISL_291186 | A/Common_Teal/Netherlands/12002960/2012 | H8N4 | | | | | present | | | | 1 |
| EPI_ISL_291189 | A/Swan/Netherlands/14000281/2014 | H5N2 | | | | | present | | | | 1 |
| EPI_ISL_291193 | A/Mallard/Netherlands/09014545/2009 | H4N6 | | | | | present | | | | 1 |
| EPI_ISL_291195 | A/Mallard/Netherlands/14016619/2014 | H3N6 | present | | | | | | | | 1 |
| EPI_ISL_293992 | A/chicken/France/160001/2016 | H5N2 | | | | | | | | present | 1 |
| EPI_ISL_293997 | A/chicken/France/160013/2016 | H5N2 | | | | | | | | present | 1 |
| EPI_ISL_294001 | A/duck/Mongolia/17/2011 | H4N3 | | | | | | present | | | 1 |
| EPI_ISL_294758 | A/mallard/France/150018/2015 | H5N1 | | | | | | | present | | 1 |
| EPI_ISL_294778 | A/duck/France/160056/2016 | H6N2 | | | | | | | | present | 1 |
| EPI_ISL_294779 | A/duck/France/160051/2016 | H6N2 | | | | | | | | present | 1 |
| EPI_ISL_297463 | A/chicken/Kostroma/1717/2017 | H5N2 | | | | | present | | | | 1 |
| EPI_ISL_297464 | A/chicken/Kostroma/1719/2017 | H5N2 | | | | | present | | | | 1 |
| EPI_ISL_297465 | A/chicken/Kostroma/1720/2017 | H5N2 | | | | | present | | | | 1 |
| EPI_ISL_297466 | A/chicken/Kostroma/1721/2017 | H5N2 | | | | | present | | | | 1 |
| EPI_ISL_298543 | A/duck/Mongolia/62/2013 | H3N1 | | | | | | present | | | 1 |
| EPI_ISL_300529 | A/teal/Markeev/13-3-11/2016 | H5N2 | | | | | | present | | | 1 |
| EPI_ISL_3007 | A/duck/Spain/539/2006 | H6N8 | | present | | | | | | | 1 |
| EPI_ISL_3008 | A/duck/Spain/543/2006 | H6N8 | | present | | | | | | | 1 |
| EPI_ISL_301004 | A/wildbird/Ibaraki/R-5W-12/2010 | H4N6 | | | | | | present | | | 1 |
| EPI_ISL_30793 | A/mallard/Netherlands/3/2005 | H3N8 | | present | | | | | | | 1 |
| EPI_ISL_30804 | A/common_eider/Netherlands/1/2006 | H3N8 | | present | | | | | | | 1 |
| EPI_ISL_30808 | A/common_teal/Netherlands/2/2005 | H6N8 | | present | | | | | | | 1 |
| EPI_ISL_30809 | A/black-headed_gull/Netherlands/1/2005 | H6N8 | | present | | | | | | | 1 |
| EPI_ISL_30813 | A/mallard/Netherlands/33/2006 | H7N8 | | present | | | | | | | 1 |
| EPI_ISL_309804 | A/Chicken/Netherlands/11004004/2011 | H8N4 | | | | | present | | | | 1 |
| EPI_ISL_309809 | A/Chicken/Netherlands/12002495-001-005/2012 | H10N9 | | | | present | | | | | 1 |
| EPI_ISL_309825 | A/Turkey/Netherlands/12004763-001-004/2012 | H10N9 | | | | present | | | | | 1 |
| EPI_ISL_309826 | A/Turkey/Netherlands/12005615/2012 | H10N9 | | | | present | | | | | 1 |
| EPI_ISL_309832 | A/Duck/Netherlands/06027358/2006 | H3N8 | present | | | | | | | | 1 |
| EPI_ISL_309833 | A/Duck/Netherlands/14015610/2014 | H6N2 | | | | present | | | | | 1 |
| EPI_ISL_309835 | A/Duck/Netherlands/14016396/2014 | H6N2 | | | | present | | | | | 1 |
| EPI_ISL_309930 | A/Chicken/Netherlands/14016059/2014 | H6N2 | | | | present | | | | | 1 |
| EPI_ISL_31484 | A/mallard/Hungary/19616/2007 | H3N8 | | present | | | | | | | 1 |
| EPI_ISL_32005 | A/mute_swan/Germany/R2927/2007 | H6N8 | | present | | | | | | | 1 |
| EPI_ISL_327464 | A/Duck/Mongolia/751/2017 | H7N3 | | | | | | | present | | 1 |
| EPI_ISL_328941 | A/ruddy_turnstone/Netherlands/2/2009 | H10N4 | | | | | | | present | | 1 |
| EPI_ISL_328956 | A/mallard_duck/Netherlands/59/2015 | H6N5 | | | | present | | | | | 1 |
| EPI_ISL_328957 | A/mallard_duck/Netherlands/60/2015 | H6N5 | | | | present | | | | | 1 |
| EPI_ISL_328962 | A/mallard_duck/Netherlands/35/2015 | H4N6 | | | | | present | | | | 1 |
| EPI_ISL_328964 | A/mallard_duck/Netherlands/11/2015 | unknown | | | | present | | | | | 1 |
| EPI_ISL_328975 | A/mallard_duck/Georgia/2/2016 | H7N7 | | | | | | present | | | 1 |
| EPI_ISL_328976 | A/mallard_duck/Georgia/10/2016 | H7N7 | | | | | | | present | | 1 |
| EPI_ISL_328985 | A/ruddy_turnstone/Netherlands/1/2009 | H10N4 | | | | | | | present | | 1 |
| EPI_ISL_328993 | A/mallard_duck/Netherlands/6/2008 | H3N6 | present | | | | | | | | 1 |
| EPI_ISL_329002 | A/mallard_duck/Georgia/1/2014 | H10N7 | | | | | | present | | | 1 |
| EPI_ISL_329023 | A/ruddy_turnstone/Netherlands/6/2008 | H3N8 | | present | | | | | | | 1 |
| EPI_ISL_329027 | A/mallard_duck/Netherlands/6/2009 | H7N7 | | | | | | | present | | 1 |
| EPI_ISL_329031 | A/mallard_duck/Netherlands/62/2008 | H3N6 | present | | | | | | | | 1 |
| EPI_ISL_329032 | A/mallard_duck/Netherlands/4/2009 | H7N7 | | | | | | | present | | 1 |
| EPI_ISL_329054 | A/White-fronted_Goose/Netherlands/3/2008 | H6N8 | | present | | | | | | | 1 |
| EPI_ISL_329056 | A/Eurasian_wigeon/Netherlands/6/2005 | H6N2 | | | | present | | | | | 1 |
| EPI_ISL_333611 | A/common_teal/Chany/889/2018 | H3N6 | | | | | present | | | | 1 |
| EPI_ISL_333612 | A/common_teal/Chany/891/2018 | H3N6 | | | | | present | | | | 1 |
| EPI_ISL_333613 | A/common_teal/Chany/892/2018 | H3N6 | | | | | present | | | | 1 |
| EPI_ISL_333614 | A/common_teal/Chany/893/2018 | H3N6 | | | | | present | | | | 1 |
| EPI_ISL_333615 | A/gadwall/Chany/893/2018 | H3N8 | present | | | | | | | | 1 |
| EPI_ISL_33850 | A/mallard/Netherlands/1/2007 | H3N2 | present | | | | | | | | 1 |
| EPI_ISL_63529 | A/mallard/Czech_Republic/14516/2007 | H3N8 | | present | | | | | | | 1 |
| EPI_ISL_63531 | A/mallard/Czech_Republic/14633-16/2007 | H3N8 | | | present | | | | | | 1 |
| EPI_ISL_67401 | A/Mallard/65112/03 | H3N8 | | present | | | | | | | 1 |
| EPI_ISL_70551 | A/mallard/Korea/GH170/2007 | H7N7 | | | | | | present | | | 1 |
| EPI_ISL_73363 | A/common_teal/Sweden/1/2003 | H3N3 | present | | | | | | | | 1 |
| EPI_ISL_73371 | A/mallard/Netherlands/5/2001 | H3N6 | present | | | | | | | | 1 |
| EPI_ISL_73380 | A/mallard/Sweden/48/2002 | H11N9 | | | present | | | | | | 1 |
| EPI_ISL_74065 | A/Teal/Norway/10_1575/2007 | H3N8 | | present | | | | | | | 1 |
| EPI_ISL_79642 | A/mallard/Germany-BW/SR871/2008 | H3N8 | present | | | | | | | | 1 |
| EPI_ISL_79643 | A/mallard/Germany-BW/SR872/2008 | H3N8 | present | | | | | | | | 1 |
| EPI_ISL_79645 | A/mallard/Germany-RP/R2322/07 | H3N8 | present | | | | | | | | 1 |
| EPI_ISL_84553 | A/mallard/Netherlands/28/2006 | H3N1 | present | | | | | | | | 1 |
| EPI_ISL_84554 | A/common_eider/Netherlands2/2006 | H4N8 | | present | | | | | | | 1 |
| EPI_ISL_84560 | A/mallard/Netherlands/20/2005 | H12N8 | | present | | | | | | | 1 |
| EPI_ISL_85900 | A/duck/Italy/6207/08 | H3N6 | present | | | | | | | | 1 |
| EPI_ISL_87299 | A/shoveler/Koymor/118/2009 | H3 | | | | | present | | | | 1 |
| EPI_ISL_89979 | A/mallard/Czech_Republic/13438-29K/2010 | H11N9 | | | | | | present | | | 1 |
| EPI_ISL_89980 | A/mallard/Czech_Republic/13577-24K/2010 | H3N8 | present | | | | | | | | 1 |
| EPI_ISL_95121 | A/Anas_platyrhynchos/Camargue/091863/09 | H10N7 | | | present | | | | | | 1 |
| EPI_ISL_97364 | A/goose/Germany/R1767/2007 | H6N8 | | present | | | | | | | 1 |
