## Supplementary material for "Detection of H3N8 influenza A virus with multiple mammalian-adaptive mutations in a rescued Grey seal (*Halichoerus grypus*) pup": Table S2

**Table S2A. Closest related viruses to A/harbor_seal/Massachusetts/1/2011**

| Segment | Seg no. | Strain | Tree Support | Host | Location | Year | Month | Subtype | Equally close on tree |
|---|---|---|---|---|---|---|---|---|---|
| HA | 4 | EPI_ISL_137683\|A/blue-winged_teal/New_Brunswick/00291/2010\|A_/_H3N8\|2010-09-14 | 0.834 | Blue-winged_teal | New_Brunswick | 2010 | 9 | H3N8 | A/mallard/Ohio/11OS2146/2011 |
| MP | 7 | EPI_ISL_131756\|A/American_black_duck/New_Brunswick/03552/2009\|A_/_H4N6\|2009-09-11 | 0.903 | American_black_duck | New_Brunswick | 2009 | 9 | H4N6 | A/American_black_duck/New_Brunswick/00914/2010 |
| NA | 6 | EPI_ISL_141230\|A/northern_pintail/Minnesota/AI09-4322/2009\|A_/_H3N8\|2009-09-12 | 0.794 | Northern_pintail | Minnesota | 2009 | 9 | H3N8 | A/American_black_duck/New_Brunswick/03559/2009 |
| NP | 5 | EPI_ISL_131756\|A/American_black_duck/New_Brunswick/03552/2009\|A_/_H4N6\|2009-09-11 | 0.94 | American_black_duck | New_Brunswick | 2009 | 9 | H4N6 | |
| NS | 8 | EPI_ISL_131756\|A/American_black_duck/New_Brunswick/03552/2009\|A_/_H4N6\|2009-09-11 | 0.961 | American_black_duck | New_Brunswick | 2009 | 9 | H4N6 | |
| PA | 3 | EPI_ISL_131756\|A/American_black_duck/New_Brunswick/03552/2009\|A_/_H4N6\|2009-09-11 | 0.87 | American_black_duck | New_Brunswick | 2009 | 9 | H4N6 | A/mallard/Wisconsin/10OS3067/2010 |
| PB1 | 2 | EPI_ISL_131756\|A/American_black_duck/New_Brunswick/03552/2009\|A_/_H4N6\|2009-09-11 | 0.967 | American_black_duck | New_Brunswick | 2009 | 9 | H4N6 | A/American_black_duck/New_Brunswick/03511/2009; A/American_black_duck/New_Brunswick/03553/2009 |
| PB2 | 1 | EPI_ISL_131756\|A/American_black_duck/New_Brunswick/03552/2009\|A_/_H4N6\|2009-09-11 | 1 | American_black_duck | New_Brunswick | 2009 | 9 | H4N6 | A/American_black_duck/New_Brunswick/03511/2009 |

**Table S2B. Closest related viruses to A/grey_seal/England/027661/2017**

| Segment | Seg no. | Strain | Tree Support | Host | Location | Year | Month | Subtype | Equally close on tree |
|---|---|---|---|---|---|---|---|---|---|
| HA | 4 | blasthit\|EPI_ISL_267243\|A/barnacle_goose/Netherlands/2/2014\|A_/_H3N6\|2014-12-15 | 0.997 | barnacle_goose | Netherlands | 2014 | 12 | H3N6 | |
| MP | 7 | blasthit\|EPI_ISL_267386\|A/mallard_duck/Netherlands/34/2010\|A_/_H3N2\|2010-10-08 | 0.792 | mallard_duck | Netherlands | 2010 | 10 | H3N2 | |
| NA | 6 | blasthit\|EPI_ISL_129976\|A/common_gull/Norway/10_1602/2009\|A_/_H6N8\|2009-XX-XX | 0.693 | common_gull | Norway | 2009 | ? | H6N8 | blasthit\|EPI_ISL_74065\|A/Teal/Norway/10_1575/2007\|A_/_H3N8\|2007-XX-XX |
| NP | 5 | blasthit\|EPI_ISL_328964\|A/mallard_duck/Netherlands/11/2015\|A\|2015-06-05 | 0.742 | mallard_duck | Netherlands | 2015 | 6 | unknown | |
| NS | 8 | blasthit\|EPI_ISL_243531\|A/barnacle_goose/Netherlands/1/2010\|A_/_H6N8\|2010-01-08 | 0.782 | barnacle_goose | Netherlands | 2010 | 1 | H6N8 | |
| PA | 3 | blasthit\|EPI_ISL_267186\|A/mallard_duck/Netherlands/8/2014\|A_/_H7N5\|2014-10-28/ blasthit\|EPI_ISL_309813\|A/Chicken/Netherlands/13015884/2013\|A_/_H5N3\|2013-11-29 | 0.792/ 0.75 | mallard_duck/ chicken | Netherlnds | 2014/ 2013 | 10/ 11 | H7N5/ H5N3 | |
| PB1 | 2 | blasthit\|EPI_ISL_328955\|A/mallard_duck/Netherlands/52/2015\|A_/_H5N6\|2015-09-16 | 0.958 | mallard_duck | Netherlands | 2015 | 9 | H5N6 | blasthit\|EPI_ISL_328966\|A/mallard_duck/Netherlands/41/2015\|A_/_H5N1\|2015-09-15; blasthit\|EPI_ISL_294761\|A/duck/France/150213/2015\|A_/_H5N2\|2015-12-01 |
| PB2 | 1 | unresolved but within a clade | 0.917 | duck/chicken | France | 2016 | various | various | |
