## Supplementary material for "Detection of H3N8 influenza A virus with multiple mammalian-adaptive mutations in a rescued Grey seal (*Halichoerus grypus*) pup": Table S3

**Table S3A**

| Strain | # glycosylation sites | 23 | 24 | 38 | 54 | 181 | 301 | 499 |
|---|---|---|---|---|---|---|---|---|
| query\|A/grey_seal/England/027661/2017\|H3N8\|HA\|2017-02_7gs | 7 | X | X | X | X | X | X | X |
| blasthit\|EPI_ISL_267243\|A/barnacle_goose/Netherlands/2/2014\|A_/_H3N6\|2014-12-15_7gs | 7 | X | X | X | X | X | X | X |
| blasthit\|EPI_ISL_291185\|A/Mallard/Netherlands/11015220/2011\|A_/_H3N8\|2011-08-26_6gs | 6 | . | X | X | X | X | X | X |
| blasthit\|EPI_ISL_243492\|A/mallard_duck/Netherlands/7/2008\|A_/_H3N8\|2008-08-22_6gs | 6 | . | X | X | X | X | X | X |
| blasthit\|EPI_ISL_328993\|A/mallard_duck/Netherlands/6/2008\|A_/_H3N6\|2008-08-22_6gs | 6 | . | X | X | X | X | X | X |
| blasthit\|EPI_ISL_267342\|A/mallard_duck/Netherlands/2/2013\|A_/_H3N6\|2013-09-10_6gs | 6 | . | X | X | X | X | X | X |
| blasthit\|EPI_ISL_291195\|A/Mallard/Netherlands/14016619/2014\|A_/_H3N6\|2014-12-03_6gs | 6 | . | X | X | X | X | X | X |
| blasthit\|EPI_ISL_243553\|A/mallard_duck/Netherlands/6/2013\|A_/_H3N8\|2013-10-01_6gs | 6 | . | X | X | X | X | X | X |
| blasthit\|EPI_ISL_309832\|A/Duck/Netherlands/06027358/2006\|A_/_H3N8\|2006-09-27_6gs | 6 | . | X | X | X | X | X | X |
| blasthit\|EPI_ISL_267207\|A/mallard_duck/Netherlands/60/2006\|A_/_H3N8\|2006-09-18_6gs | 6 | . | X | X | X | X | X | X |
| blasthit\|EPI_ISL_257067\|A/duck/Bangladesh/26920/2015\|A_/_H3N6\|2015-12-09_6gs | 6 | . | X | X | X | X | X | X |
| blasthit\|EPI_ISL_257070\|A/duck/Bangladesh/26918/2015\|A_/_H3N6\|2015-12-09_6gs | 6 | . | X | X | X | X | X | X |
| blasthit\|EPI_ISL_240672\|A/shoveler/Chany/82K/2014\|A_/_H3N8\|2014-09-10_6gs | 6 | . | X | X | X | X | X | X |
| blasthit\|EPI_ISL_333615\|A/gadwall/Chany/893/2018\|A_/_H3N8\|2018-10-19_6gs | 6 | . | X | X | X | X | X | X |
| blasthit\|EPI_ISL_257079\|A/duck/Bangladesh/26948/2015\|A_/_H3N6\|2015-12-09_6gs | 6 | . | X | X | X | X | X | X |
| blasthit\|EPI_ISL_257089\|A/duck/Bangladesh/26974/2015\|A_/_H3N6\|2015-12-10_6gs | 6 | . | X | X | X | X | X | X |
| blasthit\|EPI_ISL_243639\|A/turnstone/Netherlands/2/2007\|A_/_H3N8\|2007-07-31_6gs | 6 | . | X | X | X | X | X | X |
| blasthit\|EPI_ISL_243402\|A/dunlin/Netherlands/2/2007\|A_/_H3N8\|2007-07-31_6gs | 6 | . | X | X | X | X | X | X |
| blasthit\|EPI_ISL_243636\|A/dunlin/Netherlands/3/2007\|A_/_H3N8\|2007-07-31_6gs | 6 | . | X | X | X | X | X | X |
| blasthit\|EPI_ISL_243606\|A/gadwall_duck/Netherlands/3/2006\|A_/_H3N8\|2006-09-03_6gs | 6 | . | X | X | X | X | X | X |
| blasthit\|EPI_ISL_85900\|A/duck/Italy/6207/08\|A_/_H3N6\|2008-11-27_6gs | 6 | . | X | X | X | X | X | X |
| blasthit\|EPI_ISL_189686\|A/common_Teal/Republic_of_Georgia/1/2011\|A_/_H3N8\|2011-09-19_6gs | 6 | . | X | X | X | X | X | X |
| blasthit\|EPI_ISL_33850\|A/mallard/Netherlands/1/2007\|A_/_H3N2\|2007-XX-XX_6gs | 6 | . | X | X | X | X | X | X |
| blasthit\|EPI_ISL_237791\|A/mallard/Germany/439/2004\|A_/_H3N2\|2004-01-01_6gs | 6 | . | X | X | X | X | X | X |
| blasthit\|EPI_ISL_120213\|A/mallard/Ukraine/05842-NAMRU3/2006\|A_/_H3N8\|2006-09-30_6gs | 6 | . | X | X | X | X | X | X |
| blasthit\|EPI_ISL_89980\|A/mallard/Czech_Republic/13577-24K/2010\|A_/_H3N8\|2010-09-16_6gs | 6 | . | X | X | X | X | X | X |
| blasthit\|EPI_ISL_79643\|A/mallard/Germany-BW/SR872/2008\|A_/_H3N8\|2008-09-08_6gs | 6 | . | X | X | X | X | X | X |
| blasthit\|EPI_ISL_79642\|A/mallard/Germany-BW/SR871/2008\|A_/_H3N8\|2008-09-08_6gs | 6 | . | X | X | X | X | X | X |
| blasthit\|EPI_ISL_84553\|A/mallard/Netherlands/28/2006\|A_/_H3N1\|2006-XX-XX_6gs | 6 | . | X | X | X | X | X | X |
| blasthit\|EPI_ISL_329031\|A/mallard_duck/Netherlands/62/2008\|A_/_H3N6\|2008-10-15_6gs | 6 | . | X | X | X | X | X | X |
| blasthit\|EPI_ISL_267386\|A/mallard_duck/Netherlands/34/2010\|A_/_H3N2\|2010-10-08_6gs | 6 | . | X | X | X | X | X | X |
| blasthit\|EPI_ISL_267383\|A/mallard_duck/Netherlands/2/2003\|A_/_H3N8\|2003-09-25_6gs | 6 | . | X | X | X | X | X | X |
| blasthit\|EPI_ISL_267251\|A/mallard_duck/Netherlands/1/2003\|A_/_H3N8\|2003-09-25_6gs | 6 | . | X | X | X | X | X | X |
| blasthit\|EPI_ISL_200447\|A/duck/Hunan/04.14_YYGK431-P/2015_Mixed_\|A_/_H3N8\|2015-04-14_6gs | 6 | . | X | X | X | X | X | X |
| blasthit\|EPI_ISL_30805\|A/turnstone/Netherlands/1/2007\|A_/_H3N8\|2007-XX-XX_6gs | 6 | . | X | X | X | X | X | X |
| blasthit\|EPI_ISL_267236\|A/mallard_duck/Netherlands/51/2008\|A_/_H3N2\|2008-10-08_6gs | 6 | . | X | X | X | X | X | X |
| blasthit\|EPI_ISL_209135\|A/duck/Mongolia/996/2015\|A_/_H3N8\|2015-08-31_5gs | 5 | . | . | X | X | X | X | X |
| blasthit\|EPI_ISL_158324\|A/mallard/Sweden/79230/2008\|A\|2008-11-13_6gs | 6 | . | X | X | X | X | X | X |
| blasthit\|EPI_ISL_79645\|A/mallard/Germany-RP/R2322/07\|A_/_H3N8\|2007-XX-XX_6gs | 6 | . | X | X | X | X | X | X |
| blasthit\|EPI_ISL_73371\|A/mallard/Netherlands/5/2001\|A_/_H3N6\|2001-10-05_6gs | 6 | . | X | X | X | X | X | X |
| blasthit\|EPI_ISL_73363\|A/common_teal/Sweden/1/2003\|A_/_H3N3\|2003-08-30_6gs | 6 | . | X | X | X | X | X | X |
| blasthit\|EPI_ISL_239584\|A/duck/Gunma/3/2016\|A_/_H3N8\|2016-12-05_6gs | 6 | . | X | X | X | X | X | X |
| blasthit\|EPI_ISL_189613\|A/mallard/Sweden/79389/2008\|A\|2008-11-15_6gs | 6 | . | X | X | X | X | X | X |
| blasthit\|EPI_ISL_239582\|A/duck/Gunma/4/2016\|A_/_H3N8\|2016-12-05_6gs | 6 | . | X | X | X | X | X | X |
| blasthit\|EPI_ISL_328958\|A/mallard_duck/Netherlands/56/2015\|A_/_H3N2\|2015-08-22_7gs | 7 | X | X | X | X | X | X | X |
| blasthit\|EPI_ISL_267401\|A/mallard_duck/Netherlands/1/2004\|A_/_H3N8\|2004-09-18_6gs | 6 | . | X | X | X | X | X | X |
| blasthit\|EPI_ISL_237128\|A/duck/Shimane/321101/2015\|A_/_H3N8\|2015-11-XX_6gs | 6 | . | X | X | X | X | X | X |
| blasthit\|EPI_ISL_209111\|A/duck/Mongolia/30/2015\|A_/_H3N8\|2015-08-29_6gs | 6 | . | X | X | X | X | X | X |
| blasthit\|EPI_ISL_149873\|A/common_shelduck/Mongolia/2076/2011\|A_/_H3N8\|2011-10-08_6gs | 6 | . | X | X | X | X | X | X |
| blasthit\|EPI_ISL_158256\|A/mallard/Sweden/51869/2006\|A\|2006-10-22_6gs | 6 | . | X | X | X | X | X | X |
| blasthit\|EPI_ISL_158261\|A/mallard/Sweden/51976/2006\|A\|2006-10-24_6gs | 6 | . | X | X | X | X | X | X |

**Table S3B**

| Strain | # glycosylation sites | 46 | 54 | 67 | 84 | 144 | 293 | 398 | 268 | 42 |
|---|---|---|---|---|---|---|---|---|---|---|
| query\|A/grey_seal/England/027661/2017\|H3N8\|NA\|2017-02_7gs | 7 | X | X | X | X | X | X | X | . | . |
| blasthit\|EPI_ISL_243386\|A/mallard_duck/Netherlands/16/2007\|A_/_H6N8\|2007-11-15_6gs | 6 | X | X | . | X | X | X | X | . | . |
| blasthit\|EPI_ISL_243582\|A/bewicks_swan/Netherlands/8/2009\|A_/_H6N8\|2009-01-04_6gs | 6 | X | X | . | X | X | X | X | . | . |
| blasthit\|EPI_ISL_243640\|A/mallard_duck/Netherlands/76/2008\|A_/_H6N8\|2008-11-27_6gs | 6 | X | X | . | X | X | X | X | . | . |
| blasthit\|EPI_ISL_243495\|A/mallard_duck/Netherlands/78/2006\|A_/_H6N8\|2006-10-21_7gs | 7 | X | X | X | X | X | X | X | . | . |
| blasthit\|EPI_ISL_129976\|A/common_gull/Norway/10_1602/2009\|A_/_H6N8\|2009-XX-XX_7gs | 7 | X | X | X | X | X | X | X | . | . |
| blasthit\|EPI_ISL_73378\|A/mallard/Sweden/45/2002\|A_/_H11N8\|2002-11-13_7gs | 7 | X | X | X | X | X | X | X | . | . |
| blasthit\|EPI_ISL_243599\|A/greater_white-fronted_goose/Netherlands/4/2006\|A_/_H6N8\|2006-01-27_7gs | 7 | X | X | X | X | X | X | X | . | . |
| blasthit\|EPI_ISL_63529\|A/mallard/Czech_Republic/14516/2007\|A_/_H3N8\|2007-09-17_7gs | 7 | X | X | X | X | X | X | X | . | . |
| blasthit\|EPI_ISL_67401\|A/Mallard/65112/03\|A_/_H3N8\|2003-XX-XX_7gs | 7 | X | X | X | X | X | X | X | . | . |
| blasthit\|EPI_ISL_243402\|A/dunlin/Netherlands/2/2007\|A_/_H3N8\|2007-07-31_7gs | 7 | X | X | X | X | X | X | X | . | . |
| blasthit\|EPI_ISL_243636\|A/dunlin/Netherlands/3/2007\|A_/_H3N8\|2007-07-31_7gs | 7 | X | X | X | X | X | X | X | . | . |
| blasthit\|EPI_ISL_243639\|A/turnstone/Netherlands/2/2007\|A_/_H3N8\|2007-07-31_7gs | 7 | X | X | X | X | X | X | X | . | . |
| blasthit\|EPI_ISL_243390\|A/mallard_duck/Netherlands/40/2006\|A_/_H6N8\|2006-09-16_8gs | 8 | X | X | X | X | X | X | X | X | . |
| blasthit\|EPI_ISL_243462\|A/greater_white-fronted_goose/Netherlands/3/2006\|A_/_H6N8\|2006-01-03_7gs | 7 | X | X | X | X | X | X | X | . | . |
| blasthit\|EPI_ISL_267359\|A/mallard_duck/Netherlands/2/2004\|A_/_H6N8\|2004-09-27_7gs | 7 | X | X | X | X | X | X | X | . | . |
| blasthit\|EPI_ISL_243633\|A/mallard_duck/Netherlands/34/2006\|A_/_H3N8\|2006-09-11_8gs | 8 | X | X | X | X | X | X | X | X | . |
| blasthit\|EPI_ISL_30808\|A/common_teal/Netherlands/2/2005\|A_/_H6N8\|2005-XX-XX_7gs | 7 | X | X | X | X | X | X | X | . | . |
| blasthit\|EPI_ISL_243540\|A/greater_white-fronted_goose/Netherlands/4/2008\|A_/_H6N8\|2008-01-22_7gs | 7 | X | X | X | X | X | X | X | . | . |
| blasthit\|EPI_ISL_243546\|A/bean_goose/Netherlands/1/2008\|A_/_H6N8\|2008-01-22_7gs | 7 | X | X | X | X | X | X | X | . | . |
| blasthit\|EPI_ISL_329023\|A/ruddy_turnstone/Netherlands/6/2008\|A_/_H3N8\|2008-11-07_6gs | 6 | X | X | . | X | X | X | X | . | . |
| blasthit\|EPI_ISL_129672\|A/duck/Germany-NW/R2185/2006\|A_/_H6N8\|2006-XX-XX_7gs | 7 | X | X | X | X | X | X | X | . | . |
| blasthit\|EPI_ISL_30809\|A/black-headed_gull/Netherlands/1/2005\|A_/_H6N8\|2005-XX-XX_7gs | 7 | X | X | X | X | X | X | X | . | . |
| blasthit\|EPI_ISL_329054\|A/White-fronted_Goose/Netherlands/3/2008\|A_/_H6N8\|2008-01-22_7gs | 7 | X | X | X | X | X | X | X | . | . |
| blasthit\|EPI_ISL_267207\|A/mallard_duck/Netherlands/60/2006\|A_/_H3N8\|2006-09-18_7gs | 7 | X | X | X | X | X | X | X | . | . |
| blasthit\|EPI_ISL_243559\|A/greater_white-fronted_goose/Netherlands/2/2008\|A_/_H6N8\|2008-01-22_7gs | 7 | X | X | X | X | X | X | X | . | . |
| blasthit\|EPI_ISL_30804\|A/common_eider/Netherlands/1/2006\|A_/_H3N8\|2006-XX-XX_6gs | 6 | X | X | . | X | X | X | X | . | . |
| blasthit\|EPI_ISL_243638\|A/turnstone/Netherlands/3/2008\|A_/_H3N8\|2008-10-31_6gs | 6 | X | X | . | X | X | X | X | . | . |
| blasthit\|EPI_ISL_243606\|A/gadwall_duck/Netherlands/3/2006\|A_/_H3N8\|2006-09-03_7gs | 7 | X | X | X | X | X | X | X | . | . |
| blasthit\|EPI_ISL_189612\|A/mallard/Sweden/1636/2002\|A_/_H3N8\|2002-12-13_7gs | 7 | X | X | X | X | X | X | X | . | . |
| blasthit\|EPI_ISL_267401\|A/mallard_duck/Netherlands/1/2004\|A_/_H3N8\|2004-09-18_7gs | 7 | X | X | X | X | X | X | X | . | . |
| blasthit\|EPI_ISL_243522\|A/mallard_duck/Netherlands/13/2012\|A_/_H3N8\|2012-10-01_7gs | 7 | X | X | X | X | X | X | X | . | . |
| blasthit\|EPI_ISL_243651\|A/turnstone/Netherlands/4/2008\|A_/_H3N8\|2008-11-01_6gs | 6 | X | X | . | X | X | X | X | . | . |
| blasthit\|EPI_ISL_30793\|A/mallard/Netherlands/3/2005\|A_/_H3N8\|2005-XX-XX_7gs | 7 | X | X | X | X | X | X | X | . | . |
| blasthit\|EPI_ISL_243492\|A/mallard_duck/Netherlands/7/2008\|A_/_H3N8\|2008-08-22_7gs | 7 | X | X | X | X | X | X | X | . | . |
| blasthit\|EPI_ISL_84560\|A/mallard/Netherlands/20/2005\|A_/_H12N8\|2005-XX-XX_7gs | 7 | X | X | X | X | X | X | X | . | . |
| blasthit\|EPI_ISL_243486\|A/turnstone/Netherlands/1/2010\|A_/_H3N8\|2010-10-09_7gs | 7 | X | X | X | X | X | X | X | . | . |
| blasthit\|EPI_ISL_74065\|A/Teal/Norway/10_1575/2007\|A_/_H3N8\|2007-XX-XX_6gs | 6 | . | X | X | X | X | X | X | . | . |
| blasthit\|EPI_ISL_32005\|A/mute_swan/Germany/R2927/2007\|A_/_H6N8\|2007-XX-XX_7gs | 7 | X | X | X | X | X | X | X | . | . |
| blasthit\|EPI_ISL_15251\|A/duck/Eastern_China/163/2002\|A_/_H6N8\|2002-XX-XX_7gs | 7 | X | X | X | X | X | X | X | . | . |
| blasthit\|EPI_ISL_30813\|A/mallard/Netherlands/33/2006\|A_/_H7N8\|2006-XX-XX_6gs | 6 | X | X | . | X | X | X | X | . | . |
| blasthit\|EPI_ISL_30805\|A/turnstone/Netherlands/1/2007\|A_/_H3N8\|2007-XX-XX_7gs | 7 | X | X | X | X | X | X | X | . | . |
| blasthit\|EPI_ISL_97364\|A/goose/Germany/R1767/2007\|A_/_H6N8\|2007-XX-XX_7gs | 7 | X | X | X | X | X | X | X | . | . |
| blasthit\|EPI_ISL_3008\|A/duck/Spain/543/2006\|A_/_H6N8\|2006-10-07_8gs | 8 | X | X | X | X | X | X | X | . | X |
| blasthit\|EPI_ISL_2852\|A/duck/Norway/1/03\|A_/_H3N8\|2003-XX-XX_7gs | 7 | X | X | X | X | X | X | X | . | . |
| blasthit\|EPI_ISL_267383\|A/mallard_duck/Netherlands/2/2003\|A_/_H3N8\|2003-09-25_7gs | 7 | X | X | X | X | X | X | X | . | . |
| blasthit\|EPI_ISL_267251\|A/mallard_duck/Netherlands/1/2003\|A_/_H3N8\|2003-09-25_7gs | 7 | X | X | X | X | X | X | X | . | . |
| blasthit\|EPI_ISL_84554\|A/common_eider/Netherlands2/2006\|A_/_H4N8\|2006-XX-XX_6gs | 6 | X | X | . | X | X | X | X | . | . |
| blasthit\|EPI_ISL_31484\|A/mallard/Hungary/19616/2007\|A_/_H3N8\|2007-XX-XX_7gs | 7 | X | X | X | X | X | X | X | . | . |
| blasthit\|EPI_ISL_3007\|A/duck/Spain/539/2006\|A_/_H6N8\|2006-10-07_7gs | 7 | X | X | X | X | X | X | X | . | . |
| blasthit\|EPI_ISL_267292\|A/white-fronted_goose/Netherlands/5/2011\|A_/_H6N8\|2011-12-14_7gs | 7 | X | X | X | X | X | X | X | . | . |
