## Supplementary material for "Detection of H3N8 influenza A virus with multiple mammalian-adaptive mutations in a rescued Grey seal (*Halichoerus grypus*) pup": Table S4

| Strain | Subtype | Host | Date | aa-at-701 | aa-at-627 |
|---|---|---|---|---|---|
| A/mink/China/01/2014 | A/H9N2 | mink | 2014-12-2 | N | E |
| A/equine/Prague/2/1956 | A/H7N7 | equine | 1956-06-1 | N | E |
| A/equine/Detroit/3/1964 | A/H7N7 | equine | 1964-XX-X | N | E |
| A/equine/Prague/1956 | A/H7N7 | equine | 1956-XX-X | N | E |
| A/equine/Prague/1/1956 | A/H7N7 | equine | 1956-XX-X | N | E |
| A/equine/Lexington/1/1966 | A/H7N7 | equine | 1966-XX-X | N | E |
| A/equine/Argentina/1/1977 | A/H7N7 | equine | 1977-XX-X | N | E |
| A/equine/Uruguay/1063/1976 | A/H7N7 | equine | 1976-XX-X | N | E |
| A/equine/Sao_Paulo/4/1976 | A/H7N7 | equine | 1976-XX-X | N | E |
| A/equine/Kentucky/1a/1975 | A/H7N7 | equine | 1975-XX-X | N | E |
| A/equine/Cornell/16/74 | A/H7N7 | equine | 1974-XX-X | N | E |
| A/equine/London/1416/1973 | A/H7N7 | equine | 1973-XX-X | N | E |
| A/equine/Prague/1/1956 | A/H7N7 | equine | 1956-XX-X | N | E |
| A/tiger/Yunnan/tig1508/2015 | A/H5N1 | tiger | 2015-08-1 | N | E |
| A/Tiger/Suphanburi/Thailand/Ti-1/04 | A/H5N1 | Tiger | 2004-XX-X | N | E |
| A/Tiger/Thailand/VSMU-1-SPB/2004 | A/H5N1 | Tiger | 2004-XX-X | N | E |
| **A/harbor_seal/New_Hampshire/179629/2011** | **A/H3N8** | **harbor_seal** | **2011-09-2** | **N** | **E** |
| **A/harbor_seal/Massachusetts/1/2011** | **A/H3N8** | **harbor_seal** | **2011-09-X** | **N** | **E** |
| A_/Equine/Sweden/SVA111128SZ0073/VIR160172/2011 | A/H3N8 | Equine | 2011-11-2 | N | E |
| A/Equine/Sweden/SVA111206SZ0085/VIR165837/2011 | A/H3N8 | Equine | 2011-12-0 | N | E |
| A/equine/Sydney/2888-8/2007 | A/H3N8 | equine | 2007-08-2 | N | E |
| A/equine/East_Lothian/2/2018 | A/H3N8 | equine | 2018-02-2 | N | E |
| A/Equine/East_Sussex/1/2015 | A/H3N8 | Equine | 2015-11-0 | N | E |
| A/Equine/Hampshire/4/2016 | A/H3N8 | Equine | 2016-06-1 | N | E |
| A/Equine/Kent/1/15 | A/H3N8 | Equine | 2015-10-0 | N | E |
| A/equine/Buckinghamshire/1/2014 | A/H3N8 | equine | 2014-11-0 | N | E |
| A/equine/Northamptonshire/1/2013 | A/H3N8 | equine | 2013-07-1 | N | E |
| A/equine/Devon/1/11 | A/H3N8 | equine | 2011-10-0 | N | E |
| A/equine/East_Renfrewshire/2/11 | A/H3N8 | equine | 2011-11-0 | N | E |
| A/equine/Shropshire/10 | A/H3N8 | equine | 2010-05-0 | N | E |
| A/equine/Yorkshire/3/09 | A/H3N8 | equine | 2009-12-0 | N | E |
| A/equine/Perthshire/3/09 | A/H3N8 | equine | 2009-12-0 | N | E |
| A/equine/Dorset/09 | A/H3N8 | equine | 2009-11-0 | N | E |
| A/equine/Lanarkshire/09 | A/H3N8 | equine | 2009-09-0 | N | E |
| A/equine/Snailwell/98 | A/H3N8 | equine | 1998-01-0 | N | E |
| A/equine/Moulton/98 | A/H3N8 | equine | 1998-01-0 | N | E |
| A/equine/Cheshire/06 | A/H3N8 | equine | 2006-03-0 | N | E |
| A/equine/Lincolnshire/06 | A/H3N8 | equine | 2006-06-0 | N | E |
| A/equine/Dubai/1/12 | A/H3N8 | equine | 2012-05-3 | N | E |
| A/equine/Rio_Grande_do_Sul/1/12 | A/H3N8 | equine | 2012-03-0 | N | E |
| A/Equine/South_Africa/4/03 | A/H3N8 | Equine | 2003-09-1 | N | E |
| A/equine/Switzerland/P112/07 | A/H3N8 | equine | 2011-11-0 | N | E |
| A/equine/Kentucky/1/11 | A/H3N8 | equine | 2011-01-0 | N | E |
| A/equine/California/1/10 | A/H3N8 | equine | 2010-01-0 | N | E |
| A/equine/Tipperary/1/2019 | A/H3N8 | equine | 2019-01-0 | N | E |
| A/equine/LKZ/9/2012 | A/H3N8 | equine | 2012-09-X | N | E |
| A/equine/Kostanay/9/2012 | A/H3N8 | equine | 2012-09-X | N | E |
| A/equine/Santiago/TT10A/2018 | A/H3N8 | equine | 2018-02-1 | N | E |
| A/equine/Concepcion/RO7C/2018 | A/H3N8 | equine | 2018-02-2 | N | E |
| A/equine/Santiago/TT3A/2018 | A/H3N8 | equine | 2018-02-1 | N | E |
| A/equine/Concepcion/RO1C/2018 | A/H3N8 | equine | 2018-02-2 | N | E |
| A/equine/Concepcion/RO4C/2018 | A/H3N8 | equine | 2018-02-2 | N | E |
| A/equine/Santiago/TT13A/2018 | A/H3N8 | equine | 2018-02-1 | N | E |
| A/equine/Concepcion/RO6C/2018 | A/H3N8 | equine | 2018-02-2 | N | E |
| A/equine/Santiago/TT9A/2018 | A/H3N8 | equine | 2018-02-1 | N | E |
| A/equine/Concepcion/RO3C/2018 | A/H3N8 | equine | 2018-02-2 | N | E |
| A/equine/Curico/RO7B/2018 | A/H3N8 | equine | 2018-02-2 | N | E |
| A/equine/Curico/RO3B/2018 | A/H3N8 | equine | 2018-02-2 | N | E |
| A/equine/Santiago/TT1A/2018 | A/H3N8 | equine | 2018-02-1 | N | E |
| A/equine/Santiago/TT5A/2018 | A/H3N8 | equine | 2018-02-1 | N | E |
| A/equine/Georgia/121362-16/2016 | A/H3N8 | equine | 2016-XX-X | N | E |
| A/equine/Florida/146609/2011 | A/H3N8 | equine | 2011-XX-X | N | E |
| A/equine/New_York/135857/2016 | A/H3N8 | equine | 2016-XX-X | N | E |
| A/equine/Oregon/78356/2012 | A/H3N8 | equine | 2012-XX-X | N | E |
| A/equine/Montana/9564-1/2015 | A/H3N8 | equine | 2015-05-0 | N | E |
| A/equine/Ankara/1/2013 | A/H3N8 | equine | 2013-07-1 | N | E |
| A/equine/Saone-et-Loire/1/2015 | A/H3N8 | equine | 2015-XX-X | N | E |
| A/equine/Gironde/1/2014 | A/H3N8 | equine | 2014-XX-X | N | E |
| A/equine/Cambremer/1/2012 | A/H3N8 | equine | 2012-XX-X | N | E |
| A/equine/Neuville-Pres-Sees/1/2011 | A/H3N8 | equine | 2011-XX-X | N | E |
| A/equine/Belfond/6-2/2009 | A/H3N8 | equine | 2009-XX-X | N | E |
| A/equine/Jouars/4/2006 | A/H3N8 | equine | 2006-XX-X | N | E |
| A/equine/Tennessee/28A/2014 | A/H3N8 | equine | 2014-10-2 | N | E |
| A/equine/Tennessee/30A/2014 | A/H3N8 | equine | 2014-10-2 | N | E |
| A/equine/Tennessee/29A/2014 | A/H3N8 | equine | 2014-10-2 | N | E |
| A/equine/Tennessee/28B/2014 | A/H3N8 | equine | 2014-10-2 | N | E |
| A/equine/Tennessee/27A/2014 | A/H3N8 | equine | 2014-10-2 | N | E |
| A/equine/Tennessee/4A/2014 | A/H3N8 | equine | 2014-10-2 | N | E |
| A/equine/Himachal_Pradesh/CMVL-YOL2/2008 | A/H3N8 | equine | 2008-06-2 | N | E |
| A/equine/Kyonggi/SA1/2011 | A/H3N8 | equine | 2011-07-0 | N | E |
| A/equine/New_Market/1/1979 | A/H3N8 | equine | 1979-XX-X | N | E |
| A/equine/New_Market/nasalwash1/1979 | A/H3N8 | equine | 1979-XX-X | N | E |
| A/equine/New_York/146066/2007 | A/H3N8 | equine | 2007-09-2 | N | E |
| A/equine/Virginia/131054-3/2005 | A/H3N8 | equine | 2005-11-0 | N | E |
| A/equine/Montana/9233/2007 | A/H3N8 | equine | 2007-11-1 | N | E |
| A/equine/Ohio/113461-3/2005 | A/H3N8 | equine | 2005-XX-X | N | E |
| A/equine/Ohio/113461-1/2005 | A/H3N8 | equine | 2005-XX-X | N | E |
| A/equine/Ohio/113461-2/2005 | A/H3N8 | equine | 2005-XX-X | N | E |
| A/equine/Texas/117793/2005 | A/H3N8 | equine | 2005-10-0 | N | E |
| A/equine/Ohio/113461-2/2005 | A/H3N8 | equine | 2005-XX-X | N | E |
| A/equine/Texas/117793/2005 | A/H3N8 | equine | 2005-10-0 | N | E |
| A/equine/Ibaraki/1/2007 | A/H3N8 | equine | 2007-08-X | N | E |
| A/equine/Yokohama/aq13/2010 | A/H3N8 | equine | 2010-XX-X | N | E |
| A/equine/Malaysia/M201/2015 | A/H3N8 | equine | 2015-08-3 | N | E |
| A/equine/Sao_Paulo/1.19/2012 | A/H3N8 | equine | 2012-04-X | N | E |
| A/equine/Ahmedabad/1/2009 | A/H3N8 | equine | 2009-04-2 | N | E |
| A/equine/Mysore/1/2008 | A/H3N8 | equine | 2008-12-1 | N | E |
| A/equine/Katra-Jammu/6/2008 | A/H3N8 | equine | 2008-06-X | N | E |
| A/equine/Guangxi/1/2008 | A/H3N8 | equine | 2008-10-2 | N | E |
| A/equine/South_Kazakhstan/236/2012 | A/H3N8 | equine | 2012-08-1 | N | E |
| A/equine/Jilin/1/1989 | A/H3N8 | equine | 1989-XX-X | N | E |
| A/equine/Heilongjiang/SS1/2013 | A/H3N8 | equine | 2013-02-X | N | E |
| A/equine/Lincolnshire/1/2007 | A/H3N8 | equine | 2007-12-1 | N | E |
| A/equine/Richmond/1/2007 | A/H3N8 | equine | 2007-08-X | N | E |
| A/equine/Xuzhou/01/2013 | A/H3N8 | equine | 2013-08-2 | N | E |
| A/equine/Heilongjiang/1/2010 | A/H3N8 | equine | 2010-04-2 | N | E |
| A/equine/Tottori/1/07 | A/H3N8 | equine | 2007-XX-X | N | E |
| A/equine/Huabei/1/2007 | A/H3N8 | equine | 2007-12-0 | N | E |
| A/equine/Kascakew/1/1978 | A/H3N8 | equine | 1978-XX-X | N | E |
| A/equine/Sussex/1/1989 | A/H3N8 | equine | 1989-XX-X | N | E |
| A/equine/Kentucky/211/1987 | A/H3N8 | equine | 1987-XX-X | N | E |
| A/equine/Kentucky/1/1981 | A/H3N8 | equine | 1981-XX-X | N | E |
| A/equine/Newmarket/2/1993 | A/H3N8 | equine | 1993-XX-X | N | E |
| A/equine/Newmarket/1/1993 | A/H3N8 | equine | 1993-XX-X | N | E |
| A/equine/Newmarket/5/2003 | A/H3N8 | equine | 2003-XX-X | N | E |
| A/equine/Xinjiang/5/2007 | A/H3N8 | equine | 2007-XX-X | N | E |
| A/equine/Xinjiang/4/2007 | A/H3N8 | equine | 2007-XX-X | N | E |
| A/equine/Xinjiang/3/2007 | A/H3N8 | equine | 2007-XX-X | N | E |
| A/equine/Xinjiang/2/2007 | A/H3N8 | equine | 2007-XX-X | N | E |
| A/equine/Xinjiang/1/2007 | A/H3N8 | equine | 2007-XX-X | N | E |
| A/equine/Qinghai/1/1994 | A/H3N8 | equine | 1994-XX-X | N | E |
| A/equine/Inner_Mongolia/8/2008 | A/H3N8 | equine | 2008-XX-X | N | E |
| A/equine/Liaoning/9/2008 | A/H3N8 | equine | 2008-XX-X | N | E |
| A/equine/Heilongjiang/10/2008 | A/H3N8 | equine | 2008-XX-X | N | E |
| A/equine/Hubei/6/2008 | A/H3N8 | equine | 2008-XX-X | N | E |
| A/equine/Gansu/7/2008 | A/H3N8 | equine | 2008-XX-X | N | E |
| A/equine/Sachiyama/1/1971 | A/H3N8 | equine | 1971-XX-X | N | E |
| A/equine/France/1/1967 | A/H3N8 | equine | 1967-XX-X | N | E |
| A/equine/Switzerland/2225/1979 | A/H3N8 | equine | 1979-XX-X | N | E |
| A/equine/Johannesburg/1/1986 | A/H3N8 | equine | 1986-XX-X | N | E |
| A/equine/Algiers/1/1972 | A/H3N8 | equine | 1972-XX-X | N | E |
| A/equine/Kentucky/2/1980 | A/H3N8 | equine | 1980-XX-X | N | E |
| A/equine/Sussex/93753/89 | A/H3N8 | equine | 1989-XX-X | N | E |
| A/equine/Sussex/1/1989 | A/H3N8 | equine | 1989-XX-X | N | E |
| A/equine/Uruguay/1/1963 | A/H3N8 | equine | 1963-XX-X | N | E |
| A/equine/Berlin/1/1989 | A/H3N8 | equine | 1989-XX-X | N | E |
| A/equine/Fontainbleu/1/1979 | A/H3N8 | equine | 1979-XX-X | N | E |
| A/equine/Sao_Paulo/1/1969 | A/H3N8 | equine | 1969-XX-X | N | E |
| A/equine/Romania/1/1980 | A/H3N8 | equine | 1980-XX-X | N | E |
| A/equine/Switzerland/1118/1979 | A/H3N8 | equine | 1979-XX-X | N | E |
| A/equine/Italy/1062/1991 | A/H3N8 | equine | 1991-XX-X | N | E |
| A/equine/Roma/5/1991 | A/H3N8 | equine | 1991-XX-X | N | E |
| A/equine/Switzerland/173/1993 | A/H3N8 | equine | 1993-XX-X | N | E |
| A/equine/Austria/421/1992 | A/H3N8 | equine | 1992-XX-X | N | E |
| A/equine/Italy/1199/1992 | A/H3N8 | equine | 1992-XX-X | N | E |
| A/equine/Italy/824/1991 | A/H3N8 | equine | 1991-XX-X | N | E |
| A/equine/Santa_Fe/1/1985 | A/H3N8 | equine | 1985-XX-X | N | E |
| A/equine/Cordoba/18/1985 | A/H3N8 | equine | 1985-XX-X | N | E |
| A/equine/Sao_Paulo/6/1963 | A/H3N8 | equine | 1963-XX-X | N | E |
| A/equine/Kentucky/3/1986 | A/H3N8 | equine | 1986-XX-X | N | E |
| A/equine/Kentucky/2/1986 | A/H3N8 | equine | 1986-XX-X | N | E |
| A/equine/Kentucky/2/1987 | A/H3N8 | equine | 1987-XX-X | N | E |
| A/equine/California/8560/2002 | A/H3N8 | equine | 2002-XX-X | N | E |
| A/equine/California/4537/1997 | A/H3N8 | equine | 1997-XX-X | N | E |
| A/equine/Kentucky/Rosie100/1981 | A/H3N8 | equine | 1981-XX-X | N | E |
| A/equine/New_York/1/1983 | A/H3N8 | equine | 1983-XX-X | N | E |
| A/equine/Kentucky/8/1994 | A/H3N8 | equine | 1994-XX-X | N | E |
| A/equine/Kentucky/1/1991 | A/H3N8 | equine | 1991-XX-X | N | E |
| A/equine/Tennessee/5/1986 | A/H3N8 | equine | 1986-XX-X | N | E |
| A/equine/Alaska/29759/1991 | A/H3N8 | equine | 1991-XX-X | N | E |
| A/equine/Kentucky/1/1992 | A/H3N8 | equine | 1992-XX-X | N | E |
| A/equine/Texas/39655/1991 | A/H3N8 | equine | 1991-XX-X | N | E |
| A/equine/Kentucky/1277/1990 | A/H3N8 | equine | 1990-XX-X | N | E |
| A/equine/Kentucky/698/1988 | A/H3N8 | equine | 1988-XX-X | N | E |
| A/equine/Kentucky/694/1988 | A/H3N8 | equine | 1988-XX-X | N | E |
| A/equine/Kentucky/692/1988 | A/H3N8 | equine | 1988-XX-X | N | E |
| A/equine/Kentucky/1/1987 | A/H3N8 | equine | 1987-XX-X | N | E |
| A/equine/Kentucky/1/1986 | A/H3N8 | equine | 1986-XX-X | N | E |
| A/equine/California/83/1982 | A/H3N8 | equine | 1982-XX-X | N | E |
| A/equine/Kentucky/3/1981 | A/H3N8 | equine | 1981-XX-X | N | E |
| A/equine/Kentucky/2/1986 | A/H3N8 | equine | 1986-XX-X | N | E |
| A/equine/Wisconsin/1/03 | A/H3N8 | equine | 2003-XX-X | N | E |
| A/equine/California/191/2003 | A/H3N8 | equine | 2003-XX-X | N | E |
| A/equine/Massachussetts/213/2003 | A/H3N8 | equine | 2003-XX-X | N | E |
| A/equine/New_York/452/2003 | A/H3N8 | equine | 2003-XX-X | N | E |
| A/equine/New_York/1/1999 | A/H3N8 | equine | 1999-XX-X | N | E |
| A/equine/Kentucky/5/2002 | A/H3N8 | equine | 2002-XX-X | N | E |
| A/equine/Ohio/1/2003 | A/H3N8 | equine | 2003-XX-X | N | E |
| A/equine/New_York/VR-297/1983 | A/H3N8 | equine | 1983-XX-X | N | E |
| A/equine/California/103/1982 | A/H3N8 | equine | 1982-XX-X | N | E |
| A/equine/Kentucky/magnificent_genius1/1981 | A/H3N8 | equine | 1981-XX-X | N | E |
| A/equine/Georgia/13/1981 | A/H3N8 | equine | 1981-XX-X | N | E |
| A/equine/Georgia/10/1981 | A/H3N8 | equine | 1981-XX-X | N | E |
| A/equine/Georgia/9/1981 | A/H3N8 | equine | 1981-XX-X | N | E |
| A/equine/Georgia/3/1981 | A/H3N8 | equine | 1981-XX-X | N | E |
| A/equine/Georgia/1/1981 | A/H3N8 | equine | 1981-XX-X | N | E |
| A/equine/Kentucky/bitter_boredom5/1976 | A/H3N8 | equine | 1976-XX-X | N | E |
| A/equine/Kentucky/pass_the_pepper1/1976 | A/H3N8 | equine | 1976-XX-X | N | E |
| A/equine/Miami/1/1963 | A/H3N8 | equine | 1963-XX-X | N | E |
| A/equine/Kentucky/2/1981 | A/H3N8 | equine | 1981-XX-X | N | E |
| A/equine/California/1/1980 | A/H3N8 | equine | 1980-XX-X | N | E |
| A/equine/Kentucky/4/1980 | A/H3N8 | equine | 1980-XX-X | N | E |
| A/equine/Kentucky/1/1978 | A/H3N8 | equine | 1978-XX-X | N | E |
| A/equine/Idaho/37875/1991 | A/H3N8 | equine | 1991-XX-X | N | E |
| A/equine/Kentucky/5/02 | A/H3N8 | equine | 2002-XX-X | N | E |
| A/donkey/Shandong/1/2017 | A/H3N8 | donkey | 2017-03-2 | N | E |
| A/canine/Vermont/278213/2013 | A/H3N8 | canine | 2013-XX-X | N | E |
| A/canine/New_York/159903/2012 | A/H3N8 | canine | 2012-XX-X | N | E |
| A/canine/Florida/15592.1/2004 | A/H3N8 | canine | 2004-XX-X | N | E |
| A/canine/New_York/3699/2010 | A/H3N8 | canine | 2010-XX-X | N | E |
| A/canine/Pennsylvania/96978/2009 | A/H3N8 | canine | 2009-XX-X | N | E |
| A/canine/Maine/058124/2016 | A/H3N8 | canine | 2016-XX-X | N | E |
| A/canine/New_York/1623.1/2010 | A/H3N8 | canine | 2010-XX-X | N | E |
| A/canine/Connecticut/85863/2011 | A/H3N8 | canine | 2011-XX-X | N | E |
| A/canine/PA/33225.4/2010 | A/H3N8 | canine | 2010-XX-X | N | E |
| A/canine/NY/dog3c15/2009 | A/H3N8 | canine | 2009-03-X | N | E |
| A/canine/NY/dog4c06/2009 | A/H3N8 | canine | 2009-03-X | N | E |
| A/canine/NY/dog4c07/2009 | A/H3N8 | canine | 2009-03-X | N | E |
| A/canine/New_York/158402-1/2008 | A/H3N8 | canine | 2008-12-3 | N | E |
| A/canine/New_York/145353/2008 | A/H3N8 | canine | 2008-11-1 | N | E |
| A/canine/Pennsylvania/137154/2008 | A/H3N8 | canine | 2008-10-2 | N | E |
| A/canine/New_York/51854/2008 | A/H3N8 | canine | 2008-04-2 | N | E |
| A/canine/Colorado/6723-8/2008 | A/H3N8 | canine | 2008-01-1 | N | E |
| A/canine/Colorado/6723-8/2008 | A/H3N8 | canine | 2008-01-1 | N | E |
| A/canine/Pennsylvania/94930-3/2007 | A/H3N8 | canine | 2007-07-2 | N | E |
| A/canine/New_York/115719/2007 | A/H3N8 | canine | 2007-09-0 | N | E |
| A/canine/Florida/61156-2/2006 | A/H3N8 | canine | 2006-05-1 | N | E |
| A/canine/Florida/61156-2/2006 | A/H3N8 | canine | 2006-05-1 | N | E |
| A/canine/Pennsylvania/10915/2007 | A/H3N8 | canine | 2007-01-3 | N | E |
| A/canine/Pennsylvania/10909/2007 | A/H3N8 | canine | 2007-01-3 | N | E |
| A/canine/Pennsylvania/16699/2007 | A/H3N8 | canine | 2007-02-1 | N | E |
| A/canine/Kentucky/118778/2006 | A/H3N8 | canine | 2006-10-2 | N | E |
| A/canine/New_York/147926-5/2006 | A/H3N8 | canine | 2006-12-1 | N | E |
| A/canine/New_York/147926-3/2006 | A/H3N8 | canine | 2006-12-1 | N | E |
| A/canine/New_York/100528-6/2006 | A/H3N8 | canine | 2006-08-1 | N | E |
| A/canine/New_York/100528-5/2006 | A/H3N8 | canine | 2006-08-1 | N | E |
| A/canine/New_York/100528-1/2006 | A/H3N8 | canine | 2006-08-1 | N | E |
| A/canine/New_York/100525-1/2006 | A/H3N8 | canine | 2006-08-1 | N | E |
| A/canine/Florida/89911-2/2006 | A/H3N8 | canine | 2006-07-2 | N | E |
| A/canine/Florida/78592-7/2006 | A/H3N8 | canine | 2006-06-2 | N | E |
| A/canine/Florida/78592-6/2006 | A/H3N8 | canine | 2006-06-2 | N | E |
| A/canine/Florida/78592-2/2006 | A/H3N8 | canine | 2006-06-2 | N | E |
| A/canine/California/70645-4/2006 | A/H3N8 | canine | 2006-06-0 | N | E |
| A/canine/Colorado/30604/2006 | A/H3N8 | canine | 2006-03-2 | N | E |
| A/canine/Colorado/17864/2006 | A/H3N8 | canine | 2006-02-1 | N | E |
| A/canine/Colorado/17864/2006 | A/H3N8 | canine | 2006-02-1 | N | E |
| A/canine/Colorado/8880/2006 | A/H3N8 | canine | 2006-01-2 | N | E |
| A/canine/New_York/4986-2/2006 | A/H3N8 | canine | 2006-01-1 | N | E |
| A/canine/New_York/5183-6/2006 | A/H3N8 | canine | 2006-01-1 | N | E |
| A/canine/New_York/5183-6/2006 | A/H3N8 | canine | 2006-01-1 | N | E |
| A/canine/New_York/115809/2005 | A/H3N8 | canine | 2005-09-3 | N | E |
| A/canine/Massachusetts/26810/2016 | A/H3N8 | canine | 2016-02-2 | N | E |
| A/canine/NY/105447/2008 | A/H3N8 | canine | 2008-XX-X | N | E |
| A/canine/PA/111788.2/2009 | A/H3N8 | canine | 2009-XX-X | N | E |
| A/canine/VT/11039/2013 | A/H3N8 | canine | 2013-XX-X | N | E |
| A/canine/NY/120106.2/2011 | A/H3N8 | canine | 2011-XX-X | N | E |
| A/canine/Miami/2005 | A/H3N8 | canine | 2005-XX-X | N | E |
| A/canine/Jacksonville/2005 | A/H3N8 | canine | 2005-XX-X | N | E |
| A/canine/Florida/242/2003 | A/H3N8 | canine | 2003-XX-X | N | E |
| A/canine/Florida/43/2004 | A/H3N8 | canine | 2004-XX-X | N | E |
| A/camel/Mongolia/335/2012 | A/H3N8 | camel | 2012-12-0 | N | E |
| A/little_yellow-shouldered_bat/Guatemala/060/2010 | A/H17N10 | little_yellow-shouldered_bat | 2010-09-0 | N | S |
| A/little_yellow-shouldered_bat/Guatemala/164/2009 | A/H17N10 | little_yellow-shouldered_bat | 2009-05-0 | N | S |
| A/little_yellow-shouldered_bat/Guatemala/153/2009 | A/H17N10 | little_yellow-shouldered_bat | 2009-05-0 | N | S |
| A/Mink/Sweden/3900/1984 | A/H10N4 | Mink | 1984-10-X | N | E |
| A/flat-faced_bat/Peru/033/2010 | A/H0N0 | flat-faced_bat | 2010-XX-X | N | S |
| A/mink/Shandong/F10/2013 | A/H9N2 | mink | 2013-09-0 | D | K |
| A/mink/Shandong/F6/2013 | A/H9N2 | mink | 2013-08-1 | D | K |
| A/mink/China/02/2014 | A/H9N2 | mink | 2014-12-2 | D | E |
| A/ferret/Maryland/P10-UMD/2008 | A/H9N2 | ferret | 2008-XX-X | D | K |
| A/ferret/Maryland/P10-UMD_CIP046_RGJS08/2008 | A/H9N2 | ferret | 2008-XX-X | D | E |
| A/ferret/Maryland/P10-UMD_CIP046_RGJS07/2008 | A/H9N2 | ferret | 2008-XX-X | D | E |
| A/ferret/Maryland/P10-UMD/2012 | A/H9N2 | ferret | 2012-01-0 | D | E |
| A/equine/Guangxi/3/2011 | A/H9N2 | equine | 2011-03-1 | D | E |
| A/canine/Guangxi/1/2011/H9N2 | A/H9N2 | canine | 2011-XX-X | D | E |
| **A/seal/Massachusetts/1/1980** | **A/H7N7** | **seal** | **1980-XX-X** | **D** | **E** |
| A/feline/New_York/WVDL-20/2016 | A/H7N2 | feline | 2016-XX-X | D | E |
| A/feline/New_York/WVDL-16/2016 | A/H7N2 | feline | 2016-XX-X | D | E |
| A/feline/New_York/WVDL-14/2016 | A/H7N2 | feline | 2016-XX-X | D | E |
| A/feline/New_York/WVDL-9/2016 | A/H7N2 | feline | 2016-XX-X | D | E |
| A/feline/New_York/WVDL-3/2016 | A/H7N2 | feline | 2016-XX-X | D | E |
| A/feline/New_York/16-040082-1/2016 | A/H7N2 | feline | 2016-12-0 | D | E |
| A/feline/Guangdong/2/2014_H5N6_ | A/H5N6 | feline | 2015-01-0 | D | E |
| A/feline/Guangdong/1/2014_H5N6_ | A/H5N6 | feline | 2015-01-0 | D | E |
| A/feline/Guangdong/2/2015 | A/H5N6 | feline | 2015-01-1 | D | E |
| A/feline/Guangdong/1/2015 | A/H5N6 | feline | 2015-01-1 | D | E |
| A/cat/Zhejiang/C2/2016 | A/H5N6 | cat | 2016-01-1 | D | E |
| A/cat/Zhejiang/C1/2016 | A/H5N6 | cat | 2016-01-1 | D | E |
| A/cat/Sichuan/SC18/2014 | A/H5N6 | cat | 2014-05-0 | D | E |
| A/dog/Shandong/JT01/2009 | A/H5N2 | dog | 2009-11-X | D | E |
| A/tiger/Yunnan/tig1404/2014 | A/H5N1 | tiger | 2014-04-0 | D | V |
| A/tiger/Thailand/CU-T8/04 | A/H5N1 | tiger | 2004-XX-X | D | K |
| A/tiger/Thailand/CU-T6/04 | A/H5N1 | tiger | 2004-XX-X | D | K |
| A/tiger/Thailand/CU-T5/04 | A/H5N1 | tiger | 2004-XX-X | D | K |
| A/tiger/Thailand/CU-T4/04 | A/H5N1 | tiger | 2004-XX-X | D | K |
| A/tiger/Thailand/CU-T3/2004 | A/H5N1 | tiger | 2004-XX-X | D | K |
| A/tiger/Vietnam/NCVD-3797/2013 | A/H5N1 | tiger | 2013-08-0 | D | E |
| A/tiger/Yunnan/tig1412/2014 | A/H5N1 | tiger | 2014-12-1 | D | E |
| A/tiger/Jiangsu/01/2013 | A/H5N1 | tiger | 2013-01-2 | D | E |
| A/tiger/Shanghai/01/2005 | A/H5N1 | tiger | 2005-XX-X | D | E |
| A/stone_marten/Germany/R747/2006 | A/H5N1 | stone_marten | 2006-XX-X | D | K |
| A/raccoon_dog/Shandong/sd2/2005 | A/H5N1 | raccoon_dog | 2005-XX-X | D | E |
| A/raccoon_dog/Shandong/sd1/2005 | A/H5N1 | raccoon_dog | 2005-XX-X | D | E |
| A/plateau_pika/Qinghai/04/2007 | A/H5N1 | plateau_pika | 2007-04-1 | D | E |
| A/pika/Qinghai/BI/2007 | A/H5N1 | pika | 2007-04-2 | D | K |
| A/pika/Qinghai/SHK/2007 | A/H5N1 | pika | 2007-12-0 | D | E |
| A/pika/Qinghai/QW/2007 | A/H5N1 | pika | 2007-12-0 | D | E |
| A/pika/Qinghai/HMH/2007 | A/H5N1 | pika | 2007-12-0 | D | E |
| A/pika/Qinghai/GRL/2007 | A/H5N1 | pika | 2007-12-0 | D | E |
| A/Owston_s_civet/Vietnam/1/2005 | A/H5N1 | Owston_s_civet | 2005-XX-X | D | E |
| A/mink/Sweden/V907/06 | A/H5N1 | mink | 2006-03-2 | D | K |
| A/mink/China/XB/2015 | A/H5N1 | mink | 2015-06-3 | D | E |
| A/mink/China/G/2015 | A/H5N1 | mink | 2015-07-0 | D | E |
| A/lion/Hubei/1-2/2016 | A/H5N1 | lion | 2016-04-1 | D | E |
| A/Leopard/Suphanburi/Thailand/Leo-1/04 | A/H5N1 | Leopard | 2004-XX-X | D | K |
| A/ferret/Indonesia/5-F7/2005 | A/H5N1 | ferret | 2012-XX-X | D | K |
| A/ferret/Indonesia/5-F6/2005 | A/H5N1 | ferret | 2012-XX-X | D | K |
| A/ferret/Indonesia/5-F5/2005 | A/H5N1 | ferret | 2012-XX-X | D | K |
| A/ferret/Indonesia/5-F3/2005 | A/H5N1 | ferret | 2012-XX-X | D | K |
| A/ferret/Indonesia/5-F2/2005 | A/H5N1 | ferret | 2012-XX-X | D | K |
| A/ferret/Indonesia/5-F1/2005 | A/H5N1 | ferret | 2012-XX-X | D | K |
| A/feline/Indonesia/CDC1/2006 | A/H5N1 | feline | 2006-01-2 | D | E |
| A/equine/Egypt/av1/2009 | A/H5N1 | equine | 2009-03-2 | D | K |
| A/dog/Thailand-Suphanburi/KU-08/04 | A/H5N1 | dog | 2004-XX-X | D | K |
| A/civet_cat/Cambodia/X0313307/2013 | A/H5N1 | civet_cat | 2013-03-1 | D | E |
| A/cat/Germany/606/2006/ | A/H5N1 | cat | 2006-XX-X | D | K |
| A/cat/Thailand/KU-02/04 | A/H5N1 | cat | 2004-XX-X | D | K |
| A/muskrat/Buryatiya/1944/2000 | A/H4N6 | muskrat | 2000-XX-X | D | E |
| A/Caspian_seal/Russia/T1/2012 | A/H4N6 | Caspian_seal | 2012-10-0 | D | E |
| A/Caspian_seal/Russia/1884/2002 | A/H4N6 | Caspian_seal | 2002-07-2 | D | E |
| A/seal/Massachussetts/133/1982 | A/H4N5 | seal | 1982-XX-X | D | E |
| A/equine/North_Carolina/152429/2002 | A/H3N8 | equine | 2002-XX-X | D | E |
| A/canine/Zhejiang/S34/2015 | A/H3N8 | canine | 2015-01-0 | D | E |
| **A/seal/Massachusetts/3911/1992** | **A/H3N3** | **seal** | **1992-XX-X** | **D** | **E** |
| A/mink/Wisconsin/31512-7/2012 | A/H3N2 | mink | 2012-08-1 | D | E |
| A/mink/Nova_Scotia/1055488/2007 | A/H3N2 | mink | 2007-09-2 | D | E |
| A/ferret/Quebec/844/2011 | A/H3N2 | ferret | 2011-11-X | D | E |
| A/feline/Korea/FY057/2014 | A/H3N2 | feline | 2014-03-0 | D | E |
| A/feline/Guangdong/1/2012 | A/H3N2 | feline | 2012-11-X | D | E |
| A/feline/Guangdong/1/2011 | A/H3N2 | feline | 2011-08-1 | D | E |
| A/feline/Heilongjiang/ZH/2014 | A/H3N2 | feline | 2014-02-1 | D | E |
| A/feline/Korea/FY028/2010 | A/H3N2 | feline | 2010-10-X | D | E |
| A/feline/Korea/01/2010 | A/H3N2 | feline | 2010-03-X | D | E |
| A/feline/Korea/01/2010 | A/H3N2 | feline | 2010-03-X | D | E |
| A/canine/Guangxi/L2/2013 | A/H3N2 | canine | 2013-05-X | D | K |
| A/canine/Guangxi/L1/2013 | A/H3N2 | canine | 2013-05-X | D | K |
| A/canine/Illinois/145135-16-3-1/2016 | A/H3N2 | canine | 2016-XX-X | D | E |
| A/canine/Illinois/278597/2016 | A/H3N2 | canine | 2016-XX-X | D | E |
| A/canine/Georgia/101875/2015 | A/H3N2 | canine | 2015-XX-X | D | E |
| A/canine/Illinois/091375-16-1-1/2016 | A/H3N2 | canine | 2016-XX-X | D | E |
| A/canine/Illinois/077752/2016 | A/H3N2 | canine | 2016-XX-X | D | E |
| A/canine/Illinois/20170110-1A4/2016 | A/H3N2 | canine | 2016-XX-X | D | E |
| A/canine/Indiana/132276-16-1-2/2016 | A/H3N2 | canine | 2016-XX-X | D | E |
| A/canine/Illinois/41915/2015 | A/H3N2 | canine | 2015-XX-X | D | E |
| A/canine/Illinois/I0182389/2015 | A/H3N2 | canine | 2015-XX-X | D | E |
| A/canine/Indiana/1177-17-1/2017 | A/H3N2 | canine | 2017-XX-X | D | E |
| A/canine/Indiana/17031-16-2/2016 | A/H3N2 | canine | 2016-XX-X | D | E |
| A/canine/Indiana/093165-16-1-1/2016 | A/H3N2 | canine | 2016-XX-X | D | E |
| A/canine/Illinois/A37/2017 | A/H3N2 | canine | 2017-11-0 | D | E |
| A/canine/Florida/090919/2017 | A/H3N2 | canine | 2017-05-2 | D | E |
| A/canine/Pennsylvania/0261885/2015 | A/H3N2 | canine | 2015-XX-X | D | E |
| A/canine/North_Carolina/103297/2015 | A/H3N2 | canine | 2015-XX-X | D | E |
| A/canine/Texas/2100186/2015 | A/H3N2 | canine | 2015-XX-X | D | E |
| A/canine/Indiana/2093681/2015 | A/H3N2 | canine | 2015-XX-X | D | E |
| A/canine/Ohio/174509/2017 | A/H3N2 | canine | 2017-10-0 | D | E |
| A/canine/North_Carolina/090487/2017 | A/H3N2 | canine | 2017-05-2 | D | E |
| A/canine/Minnesota/139684/2017 | A/H3N2 | canine | 2017-08-1 | D | E |
| A/canine/Kentucky/174490/2017 | A/H3N2 | canine | 2017-10-0 | D | E |
| A/canine/Kentucky/128656/2017 | A/H3N2 | canine | 2017-07-2 | D | E |
| A/canine/Indiana/130636/2017 | A/H3N2 | canine | 2017-07-2 | D | E |
| A/canine/South_Carolina/132956/2017 | A/H3N2 | canine | 2017-08-0 | D | E |
| A/canine/Ohio/A97/2017 | A/H3N2 | canine | 2017-10-2 | D | E |
| A/canine/Ohio/A91/2017 | A/H3N2 | canine | 2017-10-2 | D | E |
| A/canine/Ohio/A82/2017 | A/H3N2 | canine | 2017-10-2 | D | E |
| A/canine/Guangdong/02/2014 | A/H3N2 | canine | 2014-02-X | D | E |
| A/canine/Guangdong/04/2013 | A/H3N2 | canine | 2013-11-X | D | E |
| A/canine/Guangdong/01/2014 | A/H3N2 | canine | 2014-02-X | D | E |
| A/canine/Guangdong/03/2013 | A/H3N2 | canine | 2013-11-X | D | E |
| A/canine/Florida/88807.2/2017 | A/H3N2 | canine | 2017-05-2 | D | E |
| A/canine/California/043670/2017 | A/H3N2 | canine | 2017-03-2 | D | E |
| A/canine/California/84315/2017 | A/H3N2 | canine | 2017-05-1 | D | E |
| A/canine/Georgia/89750.1/2017 | A/H3N2 | canine | 2017-05-2 | D | E |
| A/canine/Georgia/20170606-19/2017 | A/H3N2 | canine | 2017-05-2 | D | E |
| A/canine/Korea/AS-15/2013 | A/H3N2 | canine | 2013-09-1 | D | E |
| A/canine/Korea/AS-14/2013 | A/H3N2 | canine | 2013-08-0 | D | E |
| A/canine/Korea/AS-11/2013 | A/H3N2 | canine | 2013-06-0 | D | E |
| A/canine/Korea/AS-09/2012 | A/H3N2 | canine | 2012-10-1 | D | E |
| A/canine/Korea/AS-08/2012 | A/H3N2 | canine | 2012-09-1 | D | E |
| A/canine/Korea/AS-06/2012 | A/H3N2 | canine | 2012-08-2 | D | E |
| A/canine/Korea/AS-05/2012 | A/H3N2 | canine | 2012-07-1 | D | E |
| A/canine/Korea/AS-04/2012 | A/H3N2 | canine | 2012-06-1 | D | E |
| A/canine/Korea/AS-03/2012 | A/H3N2 | canine | 2012-05-1 | D | E |
| A/canine/Korea/AS-01/2009 | A/H3N2 | canine | 2009-09-1 | D | E |
| A/canine/Kentucky/20170606-23/2017 | A/H3N2 | canine | 2017-XX-X | D | E |
| A/canine/Illinois/20170110-2F1/2016 | A/H3N2 | canine | 2016-XX-X | D | E |
| A/canine/Illinois/M16-25374-14-1/2016 | A/H3N2 | canine | 2016-XX-X | D | E |
| A/canine/Illinois/M17-05782-7-1/2017 | A/H3N2 | canine | 2017-XX-X | D | E |
| A/canine/Florida/20170606-8/2017 | A/H3N2 | canine | 2017-XX-X | D | E |
| A/canine/Illinois/20170110-1G4/2016 | A/H3N2 | canine | 2016-XX-X | D | E |
| A/canine/Florida/20170606-9/2017 | A/H3N2 | canine | 2017-XX-X | D | E |
| A/canine/Illinois/M17-05782-8-1/2017 | A/H3N2 | canine | 2017-XX-X | D | E |
| A/canine/Pennsylvania/20170411-5/2017 | A/H3N2 | canine | 2017-XX-X | D | E |
| A/canine/Florida/20170606-11/2017 | A/H3N2 | canine | 2017-XX-X | D | E |
| A/canine/California/20170411-20/2017 | A/H3N2 | canine | 2017-XX-X | D | E |
| A/canine/Illinois/M16-25374-19-1/2016 | A/H3N2 | canine | 2016-XX-X | D | E |
| A/canine/Illinois/187427-16-7/2016 | A/H3N2 | canine | 2016-XX-X | D | E |
| A/canine/South_Korea/20170110-1F1/2016 | A/H3N2 | canine | 2016-XX-X | D | E |
| A/canine/Illinois/M17-05782-1-1/2017 | A/H3N2 | canine | 2017-XX-X | D | E |
| A/canine/Florida/20170606-2/2017 | A/H3N2 | canine | 2017-XX-X | D | E |
| A/canine/Illinois/M16-16391-2-1/2016 | A/H3N2 | canine | 2016-XX-X | D | E |
| A/canine/Kentucky/20170606-6/2017 | A/H3N2 | canine | 2017-XX-X | D | E |
| A/canine/Florida/20170606-17/2017 | A/H3N2 | canine | 2017-XX-X | D | E |
| A/canine/Illinois/20170110-2G1/2016 | A/H3N2 | canine | 2016-XX-X | D | E |
| A/canine/Illinois/M17-05782-4-1/2017 | A/H3N2 | canine | 2017-XX-X | D | E |
| A/canine/NorthCarolina/20170606-28/2017 | A/H3N2 | canine | 2017-XX-X | D | E |
| A/canine/Illinois/M17-05782-11-1/2017 | A/H3N2 | canine | 2017-XX-X | D | E |
| A/canine/Illinois/20170110-1G12/2016 | A/H3N2 | canine | 2016-XX-X | D | E |
| A/canine/Florida/20170606-7/2017 | A/H3N2 | canine | 2017-XX-X | D | E |
| A/canine/Illinois/M16-30480-3-1/2016 | A/H3N2 | canine | 2016-XX-X | D | E |
| A/canine/Florida/20170606-18/2017 | A/H3N2 | canine | 2017-XX-X | D | E |
| A/canine/China/LG2/2015 | A/H3N2 | canine | 2015-12-1 | D | E |
| A/canine/China/JLM2/2015 | A/H3N2 | canine | 2015-11-2 | D | E |
| A/canine/China/JLM1/2015 | A/H3N2 | canine | 2015-11-2 | D | E |
| A/canine/Illinois/077753/2016 | A/H3N2 | canine | 2016-XX-X | D | E |
| A/canine/Illinois/328292/2015 | A/H3N2 | canine | 2015-XX-X | D | E |
| A/canine/South_Korea/0173915/2015 | A/H3N2 | canine | 2015-XX-X | D | E |
| A/canine/Indiana/003018/2016 | A/H3N2 | canine | 2016-XX-X | D | E |
| A/canine/Florida/269770/2015 | A/H3N2 | canine | 2015-XX-X | D | E |
| A/canine/Illinois/283066/2015 | A/H3N2 | canine | 2015-XX-X | D | E |
| A/canine/Korea/CY053/2014 | A/H3N2 | canine | 2014-03-0 | D | E |
| A/canine/Korea/DG1/2014 | A/H3N2 | canine | 2014-02-1 | D | E |
| A/canine/Korea/BD-1/2013 | A/H3N2 | canine | 2013-05-3 | D | E |
| A/canine/Guangdong/23/2012 | A/H3N2 | canine | 2012-09-1 | D | E |
| A/canine/Guangdong/04/2011 | A/H3N2 | canine | 2011-02-X | D | E |
| A/canine/Guangdong/3/2011 | A/H3N2 | canine | 2011-03-X | D | E |
| A/canine/Guangdong/05/2011 | A/H3N2 | canine | 2011-02-X | D | E |
| A/canine/Guangdong/1/2011 | A/H3N2 | canine | 2011-02-X | D | E |
| A/canine/Thailand/CU-DC5299/2012 | A/H3N2 | canine | 2012-01-1 | D | E |
| A/canine/Liaoning/1578/2010 | A/H3N2 | canine | 2010-04-0 | D | E |
| A/canine/Beijing/253/2009 | A/H3N2 | canine | 2009-12-0 | D | E |
| A/canine/Beijing/420/2010 | A/H3N2 | canine | 2010-01-0 | D | E |
| A/canine/Beijing/295/2009 | A/H3N2 | canine | 2009-12-1 | D | E |
| A/canine/Beijing/305/2009 | A/H3N2 | canine | 2009-12-1 | D | E |
| A/canine/Beijing/359/2009 | A/H3N2 | canine | 2009-12-1 | D | E |
| A/canine/Beijing/362/2009 | A/H3N2 | canine | 2009-12-1 | D | E |
| A/canine/Beijing/364/2009 | A/H3N2 | canine | 2009-12-1 | D | E |
| A/canine/Beijing/418/2010 | A/H3N2 | canine | 2010-01-0 | D | E |
| A/canine/Beijing/511/2010 | A/H3N2 | canine | 2010-01-0 | D | E |
| A/canine/Beijing/1028/2010 | A/H3N2 | canine | 2010-02-0 | D | E |
| A/canine/Liaoning/1585/2010 | A/H3N2 | canine | 2010-04-0 | D | E |
| A/canine/Korea/KRIBB01/2011 | A/H3N2 | canine | 2011-11-0 | D | E |
| A/canine/Guangdong/2/2011 | A/H3N2 | canine | 2011-02-X | D | E |
| A/canine/Jiangsu/01/2009 | A/H3N2 | canine | 2009-12-X | D | E |
| A/canine/Jiangsu/02/2010 | A/H3N2 | canine | 2010-03-X | D | E |
| A/canine/Jiangsu/03/2010 | A/H3N2 | canine | 2010-04-X | D | E |
| A/canine/Jiangsu/04/2010 | A/H3N2 | canine | 2010-05-X | D | E |
| A/canine/Jiangsu/05/2010 | A/H3N2 | canine | 2010-05-X | D | E |
| A/canine/Jiangsu/06/2010 | A/H3N2 | canine | 2010-06-X | D | E |
| A/canine/Guangdong/1/2007 | A/H3N2 | canine | 2007-04-1 | D | E |
| A/canine/Guangdong/2/2007 | A/H3N2 | canine | 2007-10-1 | D | E |
| A/canine/Guangdong/2/2006 | A/H3N2 | canine | 2006-08-X | D | E |
| A/canine/Guangdong/1/2006 | A/H3N2 | canine | 2006-06-X | D | E |
| A/canine/China/LG1/2015 | A/H3N2 | canine | 2015-12-1 | D | E |
| A/canine/Texas/343907/2015 | A/H3N2 | canine | 2015-12-2 | D | E |
| A/canine/Wisconsin/19137/2016 | A/H3N2 | canine | 2016-02-0 | D | E |
| A/canine/Illinois/2883703/2015 | A/H3N2 | canine | 2015-05-0 | D | E |
| A/canine/Illinois/0843093/2015 | A/H3N2 | canine | 2015-03-3 | D | E |
| A/canine/Georgia/104940/2015 | A/H3N2 | canine | 2015-07-1 | D | E |
| A/canine/Korea/0589318/2015 | A/H3N2 | canine | 2015-03-0 | D | E |
| A/canine/Indiana/96198/2015 | A/H3N2 | canine | 2015-07-0 | D | E |
| A/canine/Illinois/1619144/2015 | A/H3N2 | canine | 2015-04-1 | D | E |
| A/canine/North_Carolina/109904/2015 | A/H3N2 | canine | 2015-07-2 | D | E |
| A/canine/Georgia/95391/2015 | A/H3N2 | canine | 2015-07-1 | D | E |
| A/canine/Illinois/12191/2015 | A/H3N2 | canine | 2015-04-1 | D | E |
| A/canine/Korea/S1/2012 | A/H3N2 | canine | 2012-11-0 | D | E |
| A/canine/Korea/VC123578/2012 | A/H3N2 | canine | 2012-03-1 | D | E |
| A/canine/Korea/VC378/2012 | A/H3N2 | canine | 2012-03-1 | D | E |
| A/canine/Heilongjiang/L1/2013 | A/H3N2 | canine | 2013-03-1 | D | E |
| A/canine/Liaoning/H6/2012 | A/H3N2 | canine | 2012-05-1 | D | E |
| A/canine/Liaoning/27/2012 | A/H3N2 | canine | 2012-04-1 | D | E |
| A/canine/Guangdong/12/2012 | A/H3N2 | canine | 2012-09-1 | D | E |
| A/canine/Korea/MV1/2012 | A/H3N2 | canine | 2012-11-0 | D | E |
| A/canine/Korea/CY009/2010 | A/H3N2 | canine | 2010-10-X | D | E |
| A/canine/Zhejiang/1/2010 | A/H3N2 | canine | 2010-09-3 | D | E |
| A/canine/Korea/01/2007 | A/H3N2 | canine | 2007-05-X | D | E |
| A/canine/Korea/GCVP01/2007 | A/H3N2 | canine | 2007-XX-X | D | E |
| A/canine/Korea/1/2010 | A/H3N1 | canine | 2010-02-0 | D | E |
| A/canine/Korea/VC125678/2012 | A/H3N1 | canine | 2012-03-1 | D | E |
| A/muskrat/Russia/63/2014 | A/H2N2 | muskrat | 2014-08-3 | D | E |
| A/panda/Sichuan/01-GG/2009 | A/H1N1 | panda | 2009-11-X | D | E |
| A/mink/South_Dakota/A01279427/2012 | A/H1N1 | mink | 2012-11-1 | D | E |
| A/giant_anteater/Tennessee/UTCVM07-733/2007 | A/H1N1 | giant_anteater | 2007-XX-X | D | K |
| A/ferret/Taiwan/E01/2013 | A/H1N1 | ferret | 2013-07-X | D | E |
| A/elephant_seal/California/2/2010 | A/H1N1 | elephant_seal | 2010-04-X | D | E |
| A/Elephant_seal/California/1/2010 | A/H1N1 | Elephant_seal | 2010-05-X | D | E |
| A/cheetah/California/D0912239/2009 | A/H1N1 | cheetah | 2009-11-X | D | E |
| A/cat/France/0514/2009 | A/H1N1 | cat | 2009-12-X | D | E |
| A/canine/Beijing/cau2/2009 | A/H1N1 | canine | 2009-11-0 | D | E |
| A/canine/Beijing/cau9/2009 | A/H1N1 | canine | 2009-11-2 | D | E |
| **A/Seal/Sweden/SVA0546/2014_** | **A/H10N7** | **Seal** | **2014-04-1** | **D** | **E** |
| **A/harbour_seal/Germany/1/2014** | **A/H10N7** | **harbour_seal** | **2014-10-0** | **D** | **E** |
| **A/harbour_seal/Germany/1/2014** | **A/H10N7** | **harbour_seal** | **2014-10-0** | **D** | **E** |
| **A/harbor_seal/NL/PV14-221_Ts/2015** | **A/H10N7** | **harbor_seal** | **2015-01-1** | **D** | **E** |
| **A/seal/Massachusetts/1/80** | **A/H7N7** | **seal** | **1980-XX-X** | **-** | **-** |
| A/tiger/Thailand/CU-T7/2004 | A/H5N1 | tiger | 2004-XX-X | - | K |
| A/equine/Almaty/26/2007 | A/H3N8 | equine | 2007-09-0 | - | - |
| A/equine/Athens/04/2007 | A/H3N8 | equine | 2007-05-0 | - | - |
| A/equine/Athens/02/2003 | A/H3N8 | equine | 2003-06-0 | - | - |
| A/equine/Yokohama/aq19/2009 | A/H3N8 | equine | 2009-XX-X | - | - |
| A/equine/La_Plata/1993 | A/H3N8 | equine | 1993-XX-X | - | - |
| A/equine/Hokkaido/I828/2008 | A/H3N8 | equine | 2008-XX-X | - | - |
| A/equine/Avesta/1993 | A/H3N8 | equine | 1993-XX-X | - | - |
| A/equine/Mongolia/1/2008 | A/H3N8 | equine | 2008-XX-X | - | - |
| A/equine/Kanazawa/1/2007 | A/H3N8 | equine | 2007-XX-X | - | - |
| A/canine/NY/dog23/2009 | A/H3N8 | canine | 2009-09-X | - | - |
| A/canine/NY/dog4/2009 | A/H3N8 | canine | 2009-03-X | - | - |
| A/canine/Korea/LBM412/2008 | A/H3N2 | canine | 2008-10-X | - | E |
| A/camel/Mongolia/82 | A/H1N1 | camel | 1982-XX-X | - | - |
